# Discovery of post-translationally modified self-peptides that promote hypertension

**DOI:** 10.1101/2023.07.18.549523

**Authors:** Nathaniel Bloodworth, Wei Chen, David Patrick, Amy Palubinsky, Elizabeth Phillips, Daniel Roeth, Markus Kalkum, Simon Mallal, Sean Davies, Mingfang Ao, Rocco Moretti, Jens Meiler, David G Harrison

**Affiliations:** Division of Clinical Pharmacology, Department of Medicine, Vanderbilt University Medical Center, Nashville, TN; Department of Medicine, Vanderbilt University Medical Center, Nashville, TN; Division of Infectious Diseases, Department of Medicine, Vanderbilt University Medical Center, Nashville, TN; Institute for Immunology and Infectious Diseases, Murdoch University, Murdoch, Australia; Center for Drug Safety and Immunology, Vanderbilt University Medical Center, Nashville, TN; Department of Immunology and Theranostics, Beckman Research Institute, City of Hope, Duarte, CA; Center for Structural Biology, Vanderbilt University, Nashville, TN; Department of Chemistry, Vanderbilt University, Nashville, TN; Institute for Drug Discovery, Universität Leipzig Medical School, Leipzig, Germany

## Abstract

Post translational modifications can enhance immunogenicity of self-proteins. In several conditions including hypertension, systemic lupus, and heart failure, isolevuglandins (IsoLGs) are formed by lipid peroxidation and covalently bond with protein lysine residues. Here we show that the murine class-I major histocompatibility complex (MHC-I) variant H-2D^b^ uniquely presents isoLG modified peptides and developed a computational pipeline that identifies structural features for MHC-I accommodation of such peptides. We identified isoLG-adducted peptides from renal proteins including the sodium glucose transporter 2, Cadherin 16, Kelch Domain containing protein 7A and solute carrier family 23, that are recognized by CD8^+^ T cells in tissues of hypertensive mice, induce T cell proliferation *in vitro*, and prime hypertension after adoptive transfer. Finally, we find similar patterns of isoLG-adducted antigen restriction in class-I human leukocyte antigens as in murine analogues. Thus, we have used a combined computational and experimental approach to define likely antigenic peptides in hypertension.

## Introduction

Accumulating evidence from the last several decades implicates inflammation and immune activation in the genesis of hypertension and its sequalae (1). Both the innate and adaptive immune systems contribute to hypertension associated inflammation (2), but T cell mediated responses in particular play an especially critical role. In 2007, Guzik et al demonstrated that mice deficient in mature T and B cells (RAG1^−/−^ mice) were protected from both angiotensin II and deoxycorticosterone acetate and sodium chloride (DOCA salt) induced hypertension. Adoptive transfer of T cells, but not B cells, restored the hemodynamic phenotype (3). Deletion of CD247 (CD3 ζ chain), which selectively eliminates T cells, reduced both hypertension and renal inflammation and kidney dysfunction in Dahl Salt Sensitive rats (4). Angiotensin II also induced T cell accumulation in the aortas and kidneys of mice with a humanized immune system, and increased levels of both effector memory CD4^+^ and CD8^+^ T cells that produce IFNγ and IL-17A in hypertensive humans. Likewise, CD8^+^ T cells with a senescent phenotype have been observed in the peripheral blood of hypertensive humans compared to normotensive controls (5, 6).

While both CD8^+^ and CD4^+^ T cells contribute to the hypertensive phenotype (7, 8), selective depletion of CD8^+^ T cells in mice confers partial protection against hypertension while CD4^+^ T cell depletion does not (9). Single cell sequencing performed in this same study showed that an oligoclonal population of CD8^+^ T cells, but not CD4^+^ T cells, accumulate in the kidney but not in other tissues (9). These data heavily imply the existence of antigens recognized by specific CD8^+^ T cells that are present in hypertension but not in normotensive conditions.

CD8^+^ T cell activation in hypertension is closely linked to excessive production of reactive oxygen species (ROS), resulting in the formation of electrophiles including Isolevuglandins (IsoLGs). IsoLGs are produced by free-radical mediated oxidation of arachidonic acid (10). They are highly reactive, electrophilic, short lived intermediates that rapidly form covalent bonds with free amines, especially lysine residues in proteins (11–13). IsoLG-adducted proteins accumulate in multiple inflammatory and cardiovascular diseases closely related to hypertension, including systemic lupus erythematosus (14), atrial fibrillation (15), atherosclerosis (16), and nonischemic heart failure (17). There is a marked increase in IsoLG adducts in DCs of hypertensive compared to normotensive mice, and scavenging of IsoLGs with 2-hydroxybenzylamine (2-HOBA) to inhibit adduct formation attenuates hypertension and end-organ damage in experimental hypertension. Adoptive transfer of DCs from hypertensive mice primes hypertension in recipient mice, and this is prevented if the donor mice have received 2-HOBA or if the recipient mice lack T cells. Likewise, adoptive transfer of DCs in which IsoLGs have been induced *ex vivo* primes hypertension in recipient mice. Further data in animal models show that T cells isolated from hypertensive mice proliferate when exposed to DCs presenting IsoLG modified proteins (18). The percent of monocytes containing IsoLGs is increased in hypertensive humans compared to normotensive subjects (18). Taken together, these data suggest that IsoLG-adducted peptides act as antigens for the activation of T cells in hypertension (19). We have also shown that factors common to the hypertensive milieu, including catecholamines (20), excess sodium (21, 22), and altered mechanical forces (23) increase formation of IsoLG-adducts in antigen presenting cells. Understanding the specific peptides that are IsoLG-adducted in hypertension would be extremely informative, providing insight into the cells and subcellular locations where this pathologic process occurs and provide therapeutic opportunity to intervene and arrest it.

In the present work we identify self-peptides that serve as substrates for IsoLG-adduction and CD8^+^ T cell activation. By leveraging several publicly available and custom-developed computational tools and workflows, we were able to identify a limited library of candidates which we individually tested *in vitro* and *in vivo*. We show that several of these candidate IsoLG-adducted peptides are recognized by CD8^+^ T cells, induce CD8^+^ T cell activation, and promote hypertension in mice. These studies are the first to define specific self-peptides that, when adducted by IsoLG, are responsible for T cell mediated inflammation in hypertension, carrying significant implications for the future treatment of hypertension and related illnesses.

## Results

### IsoLG-adducted peptides are H-2D^b^ restricted

The class-I major histocompatibility complex (MHC-I) displays selective peptide repertoires dictated by the amino acid composition of the antigen binding cleft (24), and can display antigens with post-translational modifications that generate an autoreactive CD8^+^ T cell response in a variety of diseases (25–30). We hypothesized that IsoLG-adducted peptides are similarly restricted in MHC-I presentation and that this restriction modulates T cell recognition. To test this, we generated two transgenic mouse strains expressing respectively truncated forms of one of two major MHC-I alleles found in C57BL/6 mice: H-2K^b^ or H-2D^b^. Each transgene was driven by a CD11c promoter and possessed both a His tag and a truncated transmembrane domain, allowing for extracellular secretion of MHC-I and its bound peptide (Supplemental Table 1, Supplemental Figure 1). Transgenic animals were treated for two weeks with angiotensin II or a sham infusion. In both strains, angiotensin II induced a similar degree of hypertension (Figure 1A). Splenocytes from these mice were then placed in culture for 3 days and the shed MHC-I adsorbed onto Ni-agarose beads (Figure 1B). The MHC-I loaded beads were then used to stimulate splenic CD8^+^ T cells from either hypertensive or sham infused wild-type mice that had been preloaded with the CellTrace CFSE proliferation marker. T cell proliferation was measured by assessing dye dilution. We found that T cells exposed to bead-bound and antigen-loaded H-2D^b^ proliferated, while those exposed to H-2K^b^ did not (Figure 1C). Furthermore, we observed T cell proliferation only when both T cells and bead-bound H-2D^b^ were isolated from hypertensive animals and not mice that had received sham infusion (Figure 1D). Treating the donor transgenic animals with the IsoLG scavenger 2-HOBA or adding to the culture a single-chain variable fragment antibody which binds all IsoLG adducts (D11) prevented T cell proliferation, suggesting that T cell activation occurred in response to hypertension-specific IsoLG-adducted antigens.

**Figure 1:**
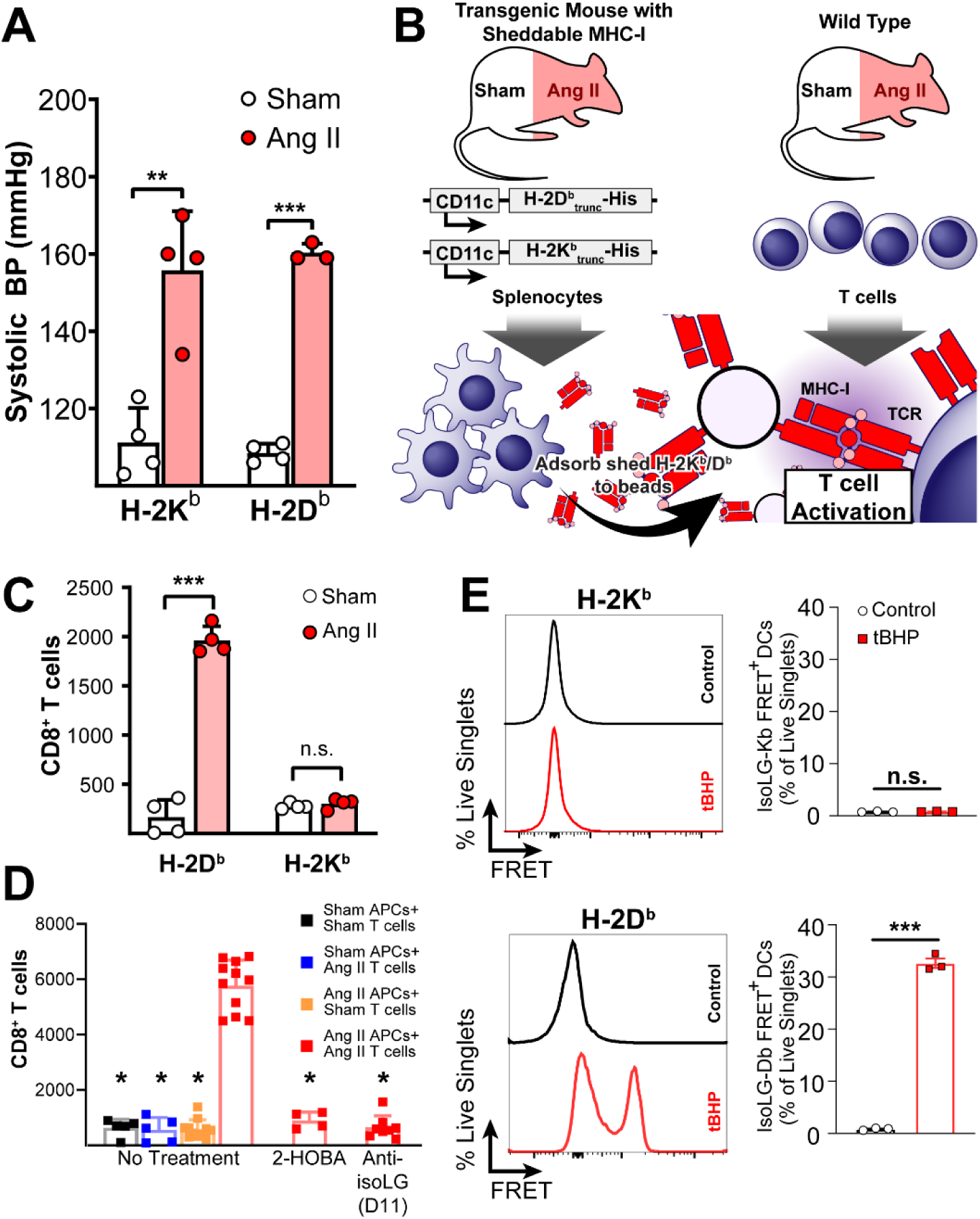
Presentation of IsoLG-adducted peptides is H-2D^b^ restricted. (A) Transgenic mice develop hypertension in response to angiotensin II (n=4, **p<0.001, \*\*\**p*<0.0001, student’s t-test). (B) Transgenic mice expressing soluble forms of H-2D^b^ or H-2K^b^ were treated with angiotensin II to induce hypertension before splenocyte harvesting and culture. Shed MHC-I was adsorbed onto Ni-agarose beads, co-cultured with T cells from wild type mice, and T cell proliferation measured with serial dye dilution and flow cytometry. (C) CD8^+^ T cells proliferate if exposed to bead-bound H-2D^b^ but not H-2K^b^ (n=4, \*\*\**p*<0.0001, student’s t-test). (D) CD8^+^ T cell proliferation is only observed if both soluble H-2D^b^ and T cells are isolated from angiotensin II-treated animals, and treating transgenic mice with the IsoLG scavenger 2-HOBA or blocking T cell/MHC-I interactions with the anti-IsoLG antibody D11 inhibits CD8^+^ T cell activation (n=4-11, *p<0.01 vs No Treatment AngII/AngII by 2-way ANOVA and Holm-Sidak post-hoc). (E) After treating mouse DCs with tBHP to induce IsoLG adduct formation, cells were stained with antibodies for MHC-I and IsoLG conjugated to a complementary FRET fluorophore pair. FRET signal was observed when staining for H-2D^b^ and IsoLG in tBHP treated DCs, but not untreated cells or when staining for H-2K^b^ (n=3, \*\*\**p*<0.0001, student’s t-test).

To determine whether IsoLG-adducted antigen restriction was due to relative differences in MHC-I binding affinity to the modified peptide between H-2D^b^ or H-2K^b^ or differences in T cell receptor recognition of the MHC-modified peptide complex, we treated murine DCs with tert-butyl hydroperoxide (tBHP), which we have shown stimulates IsoLG formation, and stained with antibodies specific for either H-2D^b^ or H-2K^b^ and IsoLG-adducted peptides. Antibodies were conjugated to complementary FRET fluorophore pairs, with a positive FRET signal indicating proximity between IsoLG-adducted peptides and either H-2D^b^ or H-2K^b^. tBHP-treated DCs stained with anti-H-2D^b^ generated a positive FRET signal while untreated cells or tBHP-treated DCs stained with H-2K^b^ did not (Figure 1E). These results indicate that H-2D^b^ can present IsoLG-adducted peptides, promoting T cell activation, while H-2K^b^ cannot.

### Computational screening identifies peptide residue positions favoring IsoLG adduction

Peptide binding affinity for MHC-I is largely dictated by structural constraints imposed by the MHC-I peptide binding cleft which, unlike the groove of MHC-II, is closed and accommodates shorter peptides 8-10 amino acids long (31–33). We reasoned that the IsoLG-adducted lysine imposed significant limitations on a peptide’s ability to take on certain structural conformations, and that understanding these limitations might help narrow the list of possible peptides serving as substrates for IsoLG adduction. To test this hypothesis, we developed a computational pipeline for modeling IsoLG-adducted peptides bound to MHC-I using FlexPepDock refinement. FlexPepDock refinement is a protocol implemented in the protein modeling software suite Rosetta, developed to model receptor-bound peptides and used previously to predict MHC-I/peptide structures and peptide-receptor binding affinity with a high degree of accuracy (34–36).

Using pre-existing structural templates, we benchmarked FlexPepDock refinement on all peptide/MHC-I structures available in the protein data bank, generating high fidelity models (Supplemental Figure 2A-C, Supplemental Table 2). We next generated structures for H-2D^b^ and H-2K^b^ bound epitopes with known binding affinity available in the immune epitope database (IEDB) and compared the Rosetta energy score terms for known binders (≤ 500nm IC_50_) and non-binders (> 500nm IC_50_) (37). For both H-2D^b^ and H-2K^b^ epitopes, known MHC-I binders had on average lower, or more favorable, Rosetta energies than non-binders (Figure 2A-B). We then selected known binders to H-2D^b^ or H-2K^b^ containing at least one lysine residue and measured changes in Rosetta energy after *in silico* IsoLG adduction. H-2K^b^ bound epitopes displayed a greater increase in Rosetta energy after IsoLG-adduction than H-2D^b^ bound epitopes (Figure 2C), suggesting an inability to accommodate this post-translational modification.

**Figure 2:**
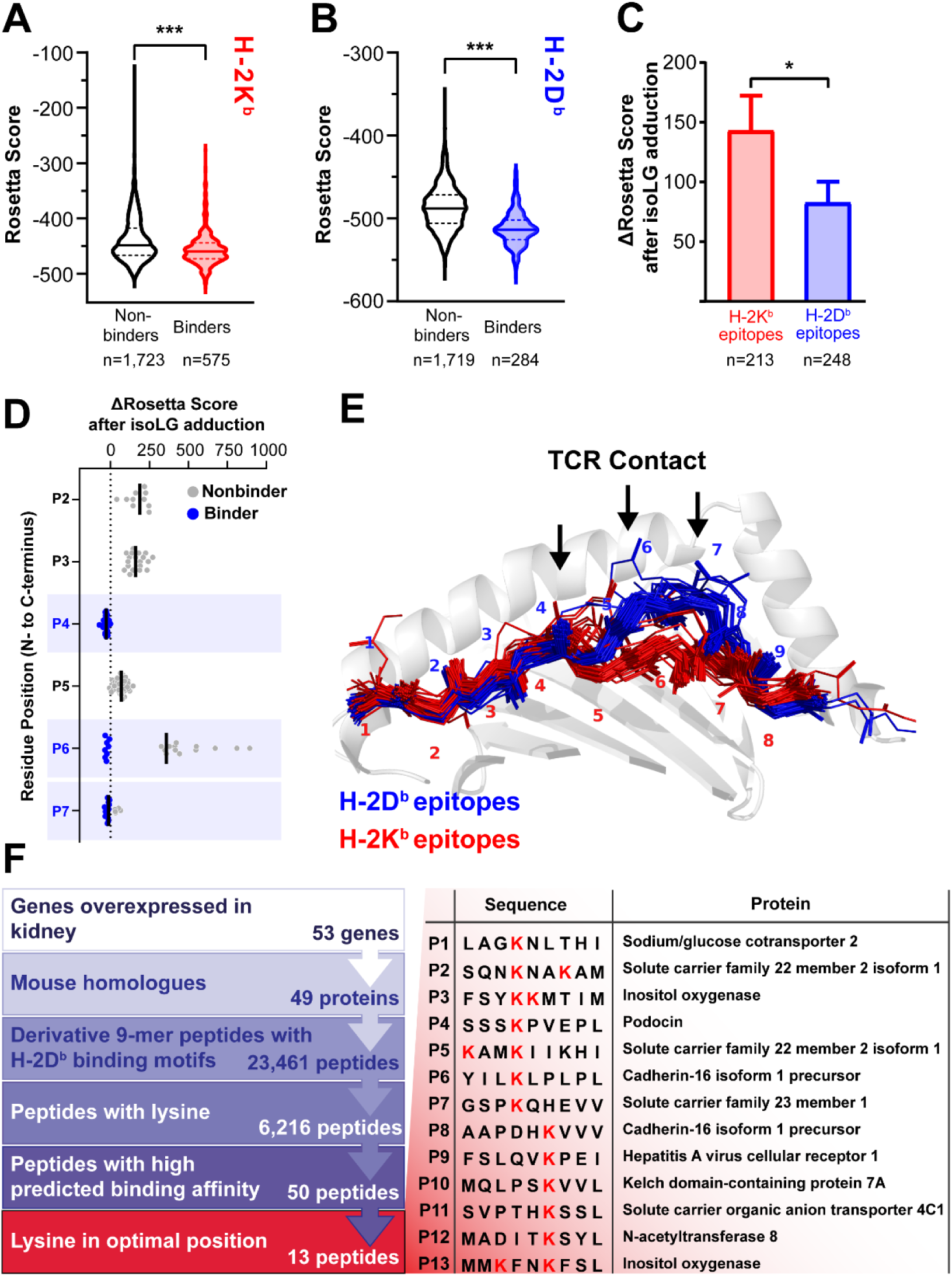
A Computational modeling pipeline predicts IsoLG adducted peptides presented by H-2D^b^. (A-B) Rosetta scores are more favorable (more negative) for peptides known to bind to H-2K^b^ (A) and H-2D^b^ (B) when compared to known non-binders (\*\*\**p*<0.0001, Mann-Whitney test). (C) When lysine-containing peptides are adducted with IsoLG *in silico*, Rosetta predicts smaller (more favorable) score changes if those peptides are bound to H-2D^b^ compared to H-2K^b^ (\**p*<0.05, Mann-Whitney test). (D) Rosetta score changes after *in silico* IsoLG adduction for all non-anchoring residues in lysine containing H-2D^b^ bound epitopes predict residue sites 4, 6, and 7 as energetically favorable positions (positions at which multiple peptides modeled have no score increase after adduction. Black bars indicate mean Δ Rosetta score). (E) These residues correspond to areas recognized by TCRs interacting with H-2D^b^ epitopes. (F) Strategy for identifying a library of peptide candidates to screen *in vitro* and *in vivo*.

We next examined per-residue energy changes for H-2D^b^ bound epitopes 8-9 residues in length before and after IsoLG-adduction for all non-anchoring residues and found peptides with lysine at residue positions 4, 6, or 7 showed the smallest changes in Rosetta energy following IsoLG adduction (Figure 2D). These positions correspond to solvent-accessible sites that are largely responsible for dictating T cell recognition in H-2D^b^ restricted epitopes (Figure 2E) (38).

Leveraging this insight, we produced a limited library of peptide candidates for further screening, each containing lysine at one of the optimal IsoLG-adduction residue positions. Given our prior evidence that the kidney is a likely source of IsoLG-adducted peptides in hypertension, we focused our initial search on proteins with relative over-expression in the kidney (39). Of the 53 candidates identified, 49 had corresponding mouse homologs. We next identified peptide sequences derived from those proteins with H-2D^b^ binding motifs, predicted by the webtool NetMHCpan 4.0 to bind to H-2D^b^ with high affinity (“strong binders” are defined by a percent rank score unique to NetMHCpan) (40). This approach yielded 13 peptides with at least one lysine in a position favoring IsoLG-adduction as predicted by Rosetta. These sequences and the proteins from which they are derived are summarized in Figure 2F and Supplemental Table 3.

### IsoLG adducted candidate peptides are recognized by T cells in hypertensive mice, stimulate T cell proliferation in vitro, and induce hypertension in vivo

We isolated T cells from the bone marrow of angiotensin II-treated mice and exposed them to DCs pulsed with each candidate peptide, measuring T cell proliferation by serial dye dilution. Seven of 13 candidate peptides induced a significant increase in proliferation of T cells from hypertensive but not normotensive mice (Figure 3A, Supplemental Figure 3). To test T cell specificity for each candidate peptide we employed a fluorescently tagged H-2D^b^/IgG1 fusion protein loaded with each candidate, IsoLG-adducted or not. Of the 13 peptides screened, 10 were identified aortic CD8^+^ T cells from hypertensive mice to a greater extent than observed in mice without hypertension (Figure 3B and Supplemental Figure 4-5). This analysis revealed that up to 14% of CD8^+^ T cells in aortas of hypertensive mice recognized individual candidate peptides that were IsoLG-adducted. T cells did not recognize peptides unadducted by IsoLGs. Six candidates induced proliferation in T cells from angiotensin II treated mice and identified CD8^+^ T cells enriched in the aortas in hypertension (Figure 3C-D).

**Figure 3:**
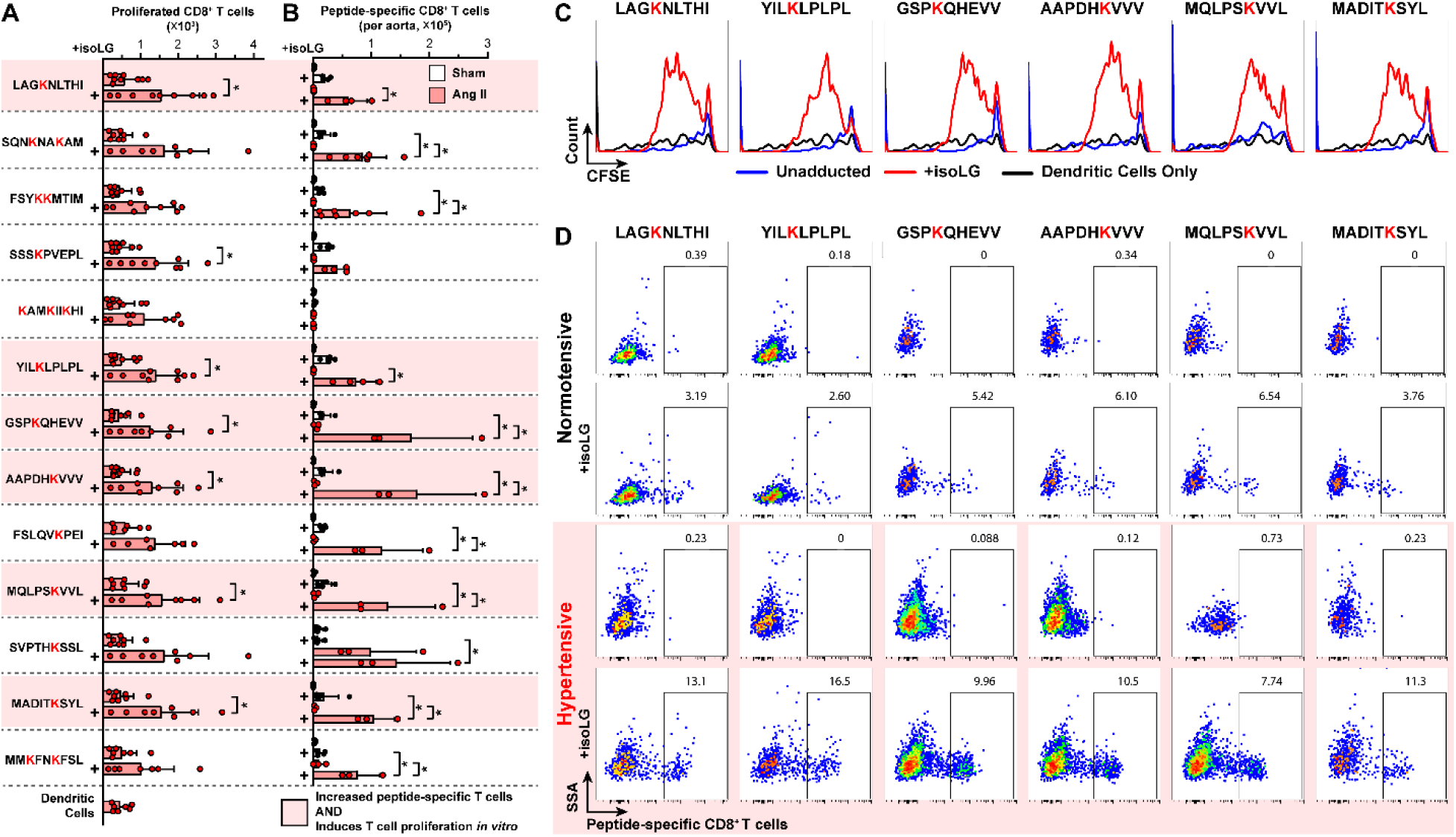
A subset of candidate IsoLG-adducted peptides are recognized by CD8^+^ T cells enriched in the aortas of hypertensive mice and induce CD8^+^ T cell prliferation *in vitro*. (A) Seven candidate IsoLG-adducted peptides induce the proliferation of T cells isolated from the bone marrow of hypertensive mice, while unadducted peptides do not (n=7-9, *p<0.05, student’s t-test). (B) 11 IsoLG-adducted peptides identify a population of peptide-specific CD8^+^ T cells that are enriched in the aortas of hypertensive mice (n=3-9,*p<0.05 by 2-way ANOVA and Holm-Sidak post-hoc). Six candidates are both recognized by CD8^+^ T cells and induce CD8^+^ T cell proliferation *in vitro* (highlighted in red). (C) Representative histograms illustrating proliferation of CD8^+^ T cells after exposure to each of these six candidate IsoLG-adducted peptides and (D) flow plots illustrating the increase in peptide-specific CD8^+^ T cells in the aorta.

Given that the bone marrow serves as a reservoir for hypertension-specific CD8^+^ memory T cells (20, 41), we also performed flow cytometry on single cell suspensions from the bone marrow of angiotensin II treated mice. CD8^+^ T cells recognizing eight of the IsoLG-adducted peptide candidates were enriched in the bone marrow of hypertensive mice compared to normotensive controls (Figure 4A). Of the six IsoLG-adducted peptides labeling CD8^+^ T cell populations enriched in the aorta, four also labeled T cells enriched in the bone marrow. Staining with the memory T cell markers CD44 and CD62L revealed that CD8^+^ T cells recognizing IsoLG-adducted peptides were predominantly effector memory cells in the aorta (Figure 4B-C), and a mixture of effector memory and central memory cells in the bone marrow (Figure 4D-E).

**Figure 4:**
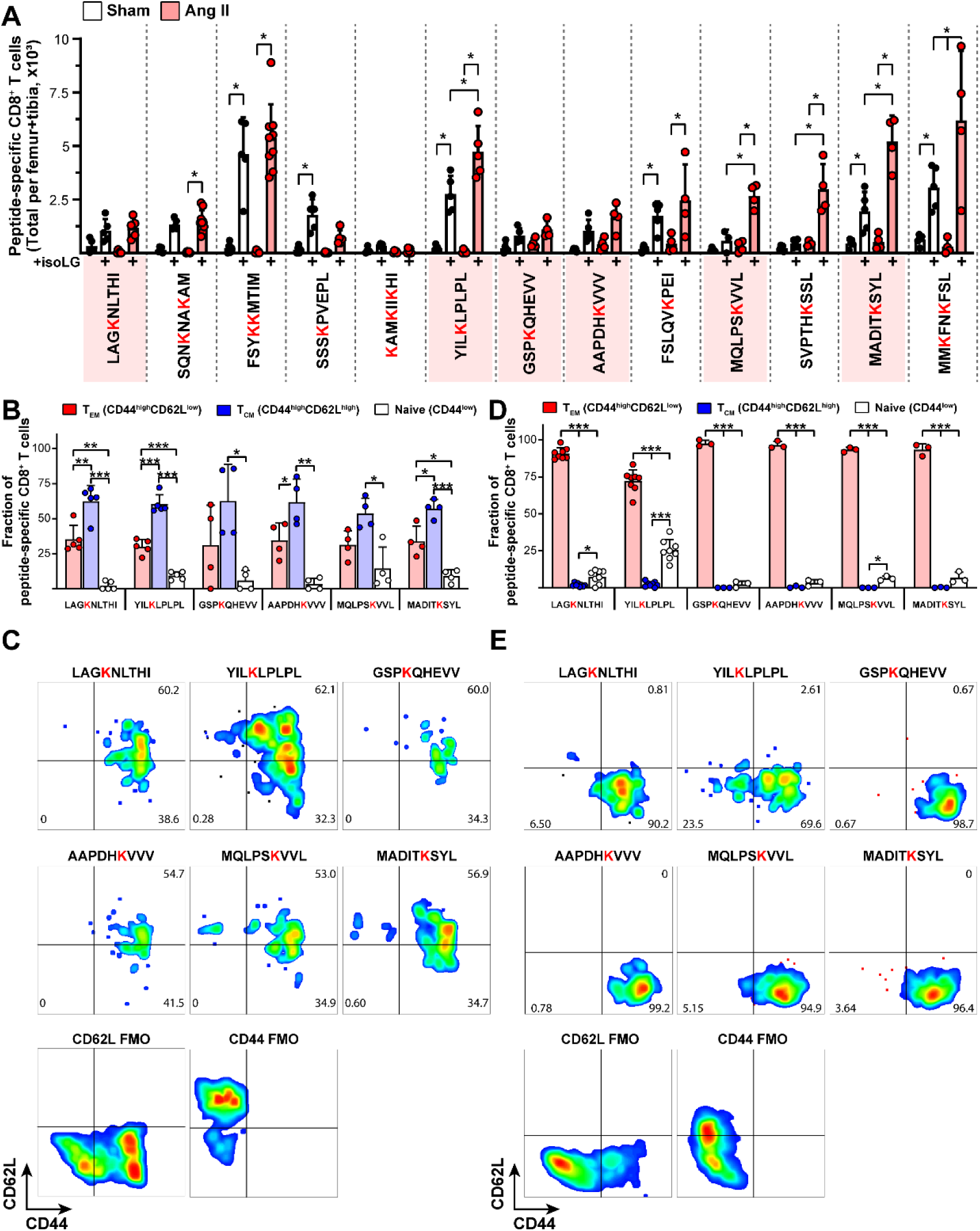
IsoLG-adducted peptide-specific CD^+^ T cells are predominantly memory T cells. (A) Candidate peptides adducted with IsoLG are recognized by CD8^+^ T cells in the bone marrow, an important reservoir for memory CD8^+^ T cells in hypertension (n=3-8, \**p*<0.05, 2-way ANOVA and Holm-Sidak post-hoc). Staining for memory T cell markers CD44 (all memory cells) and CD62L (central memory cells) reveals these populations are primarily a mix of effector and central memory cells in (B-C) the bone marrow and mostly effector memory cells in (D-E) the aorta (n=3-8, \**p*<0.05, \*\**p*<0.001, \*\*\**p*<0.0001, 1-way ANOVAs and Holm-Sidak post-hoc).

We also performed flow cytometry on kidney homogenates, staining for CD8^+^ T cells specific for the six candidate IsoLG adducted peptides that are recognized by aortic T cells and capable of inducing T cell proliferation. We confirmed that CD8^+^ T cells recognized two of the six IsoLG-adducted peptides that were enriched in the aortas of hypertensive mice, and all six were present in both sham and angiotensin II treated animals (Supplemental Figure 6A-B). These T cells were also predominantly memory effector cells like those observed in the aorta (Supplemental Figure 6C-D).

### IsoLG modified candidate peptides augment hypertension in vivo

To test the potential roles of each candidate peptide in hypertension, we performed the experiment illustrated in Figure 5A. Briefly, CD11c^+^ DCs were pulsed overnight with candidate peptides with or without IsoLG adduction. These DCs were then adoptively transferred to WT mice and five days later an infusion of a generally subpressor dose of angiotensin II was begun (18). Blood pressures were measured before and after two weeks of angiotensin II treatment. Of the six peptides tested (those that labeled CD8^+^ T cells in the aorta and induced CD8^+^ T cell proliferation *in vitro*), four induced blood pressure elevations as high as 180 mmHg after adoptive transfer (Figure 5B).

**Figure 5:**
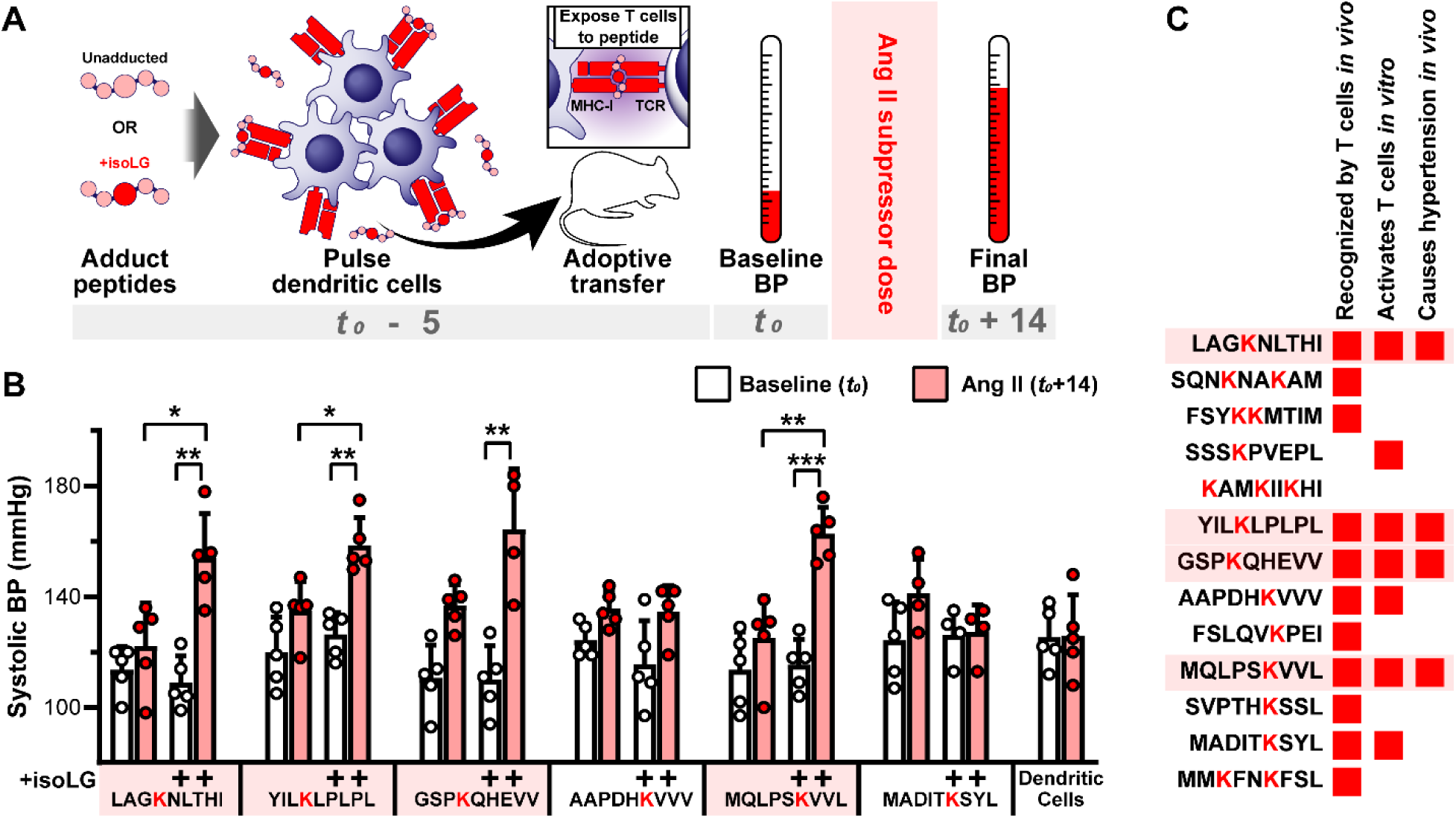
IsoLG-adducted peptides induce hypertension in mice. (A) Experimental diagram. IsoLG-adducted peptide candidates and their unadducted counterparts were loaded onto DCs and adoptively transferred prior to two weeks of treatment with a subpressor dose of angiotensin II. (B) Four of the six candidates tested induced a significant increase in blood pressure following adoptive transfer while unadducted peptides do not (n=4-5, \**p*<0.05, \*\**p*<0.001, \*\*\**p*<0.0001, 2-way ANOVA and Holm-Sidak post-hoc). (C) Of the original 13 candidates screened, four are recognized by CD8^+^ T cells, induce CD8^+^ T cell activation, and induce hypertension in mice following adoptive transfer.

Figure 5C summarizes the stepwise screening of our candidate peptides. Of the 13 peptides originally identified by our computational analysis, 11 identified T cells in the target tissues of hypertensive mice. Seven of these peptides induced proliferation of CD8^+^ T cells from hypertensive mice, and four augmented hypertension *in vivo*.

We performed mass spectrographic analysis of the un-adducted and IsoLG-adducted variants for one of the immunogenic peptides (peptide 1, LAGKNLTHI). Ions with IsoLG adducts were positively identified, but the overall signal strength was significantly lower than that of the unadducted variant (Supplemental Figure 7). Ion chromatograms demonstrated the presence of multiple diastereomers, including pyrrole, anhydrolactam, and anhydropyrrole products all present in the same sample (Supplemental Figure 8).

### Class-I human leukocyte antigens exhibit differential presentation of IsoLG-adducted peptides

The above data from mice strongly suggest that T-cell recognition of IsoLG-adducted peptides depends on their affinity for a given MHC-I. To determine if IsoLG adducted peptides are similarly restricted in their presentation among human MHC-I variants, we identified class-I human leukocyte antigen (HLA) variants from a curated database of HLA allele frequencies (42). We selected HLA alleles for each of three subpopulations from the US National Merit Donor Program with a phenotype frequency >5%, leaving 18 HLA alleles in total after excluding duplicates between subpopulations (Supplemental table 4). Cells lacking native HLA expression (HLA null) were transfected with each allele, treated with tBHP to induce IsoLG formation, and assayed for IsoLG-adduct presentation as in Figure 1E using FRET and flow cytometry. We found significant variability of FRET signal between various HLA-A and HLA-B alleles, suggesting that certain alleles are more adept at displaying IsoLG-adducted peptides (Figure 6A). Scavenging IsoLGs with ethyl-2-HOBA significantly reduced FRET signal across all HLA alleles tested (Figure 6B).

**Figure 6:**
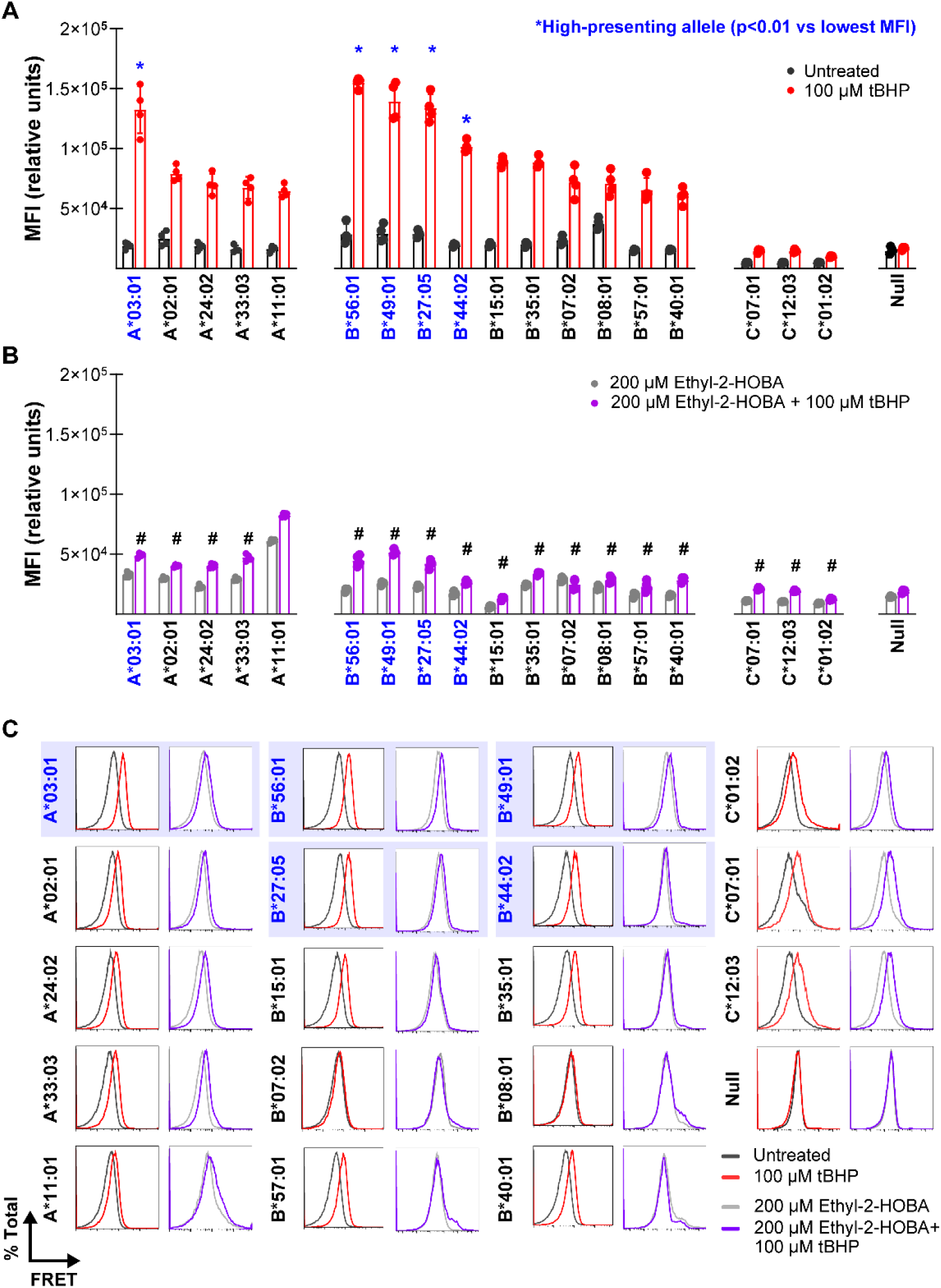
IsoLG adducted peptides are preferentially displayed by certain HLA molecules. (A) Treating K562 cells expressing single HLA alleles with tBHP induces a significant increase in the HLA-IsoLG FRET proximity mean fluorescent intensity (MFI) for all alleles screened, excepting the HLA null control. However, there are certain alleles with significantly higher FRET MFI compared to the lowest measured in each allele class (A, B, or C) (n=4, \**p*<0.05 vs lowest average MFI for corresponding allele class, 2-way ANOVA and Holm-Sidak post-hoc test). (B) Treatment with the dicarbonyl scavenger ethyl-2-HOBA significantly reduces FRET MFI compared to tBHP only treated cells (n=4, ^#^*p*<0.05 vs corresponding tBHP treated group, 2-way ANOVA with Holm-Sidak post-hoc). (C) Example FRET signal for all HLA alleles tested. “High-presenting” HLA-A and HLA-B alleles are highlighted in blue.

We identified one HLA-A variant and four HLA-B variants with an enhanced FRET signal, suggesting these alleles can display IsoLG-adducted peptides (Figure 6C). Using FlexPepDock, we selected lysine-containing peptides, nine residues in length and derived from proteins overexpressed in human renal tissue, predicted bind to the HLA in question with high affinity. We modeled the peptides to each HLA screened before and after IsoLG adduction. Rosetta energy scores were more favorable for IsoLG-adducted peptides docked to the HLA-A and HLA-B variants with enhanced FRET signal (“high-presenters”) when compared to the remaining HLA variants (Supplemental Figure 9A). For the three high-presenting HLA variants with available crystal structures, we compared energy score changes at each residue position outside of the MHC-I binding pockets and for which there was at least one lysine containing peptide. Similar to H-2D^b^ bound peptides, we found that favorable energy score changes after IsoLG adduction corresponded to more exposed sites with potentially greater T cell receptor accessibility on representative HLA-bound peptides (Supplemental Figure 9E).

## Discussion

For decades, T cells have been identified in the peripheral tissues of humans with hypertension and in several experimental models. Evidence accumulated over the last 10 years strongly implies the existence of IsoLG-adducted peptides as antigens in hypertension (9, 18, 43). In this work we identify for the first time self-peptides that, when IsoLG-modified, are both recognized by and activate CD8^+^ T cells in hypertensive mice and prime hypertension *in vivo*. These findings strongly suggest that the peptides identified in the current study or similar IsoLG-modified peptides play a role in the genesis of hypertension and its related end-organ damage.

The finding that T cells identified by these IsoLG-modified peptides are present in the aorta, kidney, and bone marrow of hypertensive mice is compatible with prior work in which we showed that an oligoclonal population of CD8^+^ T cells accumulate in the kidneys of hypertensive mice and that T cells with a memory phenotype home to the bone marrow in hypertension. We have also shown that renal denervation prevents the appearance of activated dendritic cells in secondary lymphoid organs and prevents the ultimate accumulation of T cells in the kidney. Taken together and with our current data, a paradigm emerges in which antigens in the kidney, like those identified in the current study, are presented to CD8^+^ T cells in secondary lymphoid organs. These activated T cells then home back to the kidney, vasculature, and the bone marrow, where they promote organ dysfunction and end-organ damage.

In mouse models of obesity-associated insulin resistance, oligoclonal CD8^+^ T cell populations accumulate in response to IsoLG protein adducts presented in DCs located in adipose tissue (44). Oligoclonal CD8^+^ T cell populations are also found in human and murine atherosclerotic plaques, likely in response to plaque-specific antigens (16, 45, 46). In a murine model of heart failure, CD4^+^ T cells are activated after exposure to IsoLG-adducted cardiac peptides presented by class-II MHC (17). We have also shown that IsoLG-adducted peptides contribute to systemic lupus erythematosus (14). It is unclear if the peptides identified in the present study contribute to these other diseases; however, a combined computational and experimental approach like that employed here could be useful in these related conditions.

We found that CD8^+^ T cells recognizing IsoLG-adducted peptides in the aorta, kidney, and bone marrow are both effector and central memory cells. These findings are consistent with prior experiments performed in our laboratory illustrating that memory CD8^+^ T cells accumulate in the bone marrow and kidney in response to repeated hypertensive stimuli, and that memory T cell formation through CD70 mediated co-stimulation is necessary for hypertension pathogenesis (20, 41). Similar observations have been made in humanized mice demonstrating increased numbers of memory T cells in the aorta and lymphatic tissues after induction of hypertension (5).

It is also of interest that even in non-hypertensive mice, 3 to 6% of CD8^+^ T cells in the aorta, kidney, and bone marrow recognize IsoLG-adducted peptides. In the peripheral tissues, these cells have the phenotype of effector memory cells, and in the bone marrow they are a mix of effector and central memory cells. These findings suggest that these T cells may have previously encountered molecularly similar antigens. One such source of exposure could be antigens derived from commensal organisms or other chronic persistent foreign antigens that either mimic IsoLG-adducts or that are themselves IsoLG-adducted. Hypertension is closely associated with alterations in gut microbiome content in humans (47). Germ-free mice are also protected against angiotensin II-induced hypertension and have reduced arterial and renal inflammatory infiltrate, and transplant of gut microbiome from hypertensive patients to germ free animals can induce high blood pressure (48). Salt sensitivity, an important hypertension phenotype, is also intrinsically linked to both gut microbiome alterations in mice and humans and is a potent stimulator for IsoLG production in DCs (49, 50). Whether or not the same T cells responsible for mediating inflammatory responses against gut bacteria (or some other viral antigen) also recognize IsoLG adducted antigens is an important subject of further study.

The proteins we identified from which the IsoLG adducted peptides are derived are all membrane-associated proteins and include transporters (in the case of peptide 1, derived from the sodium/glucose cotransporter encoded by Slc5a2, and P7, derived from the sodium/vitamin C cotransporter encoded by Slc23a1), cell-cell adhesion proteins (peptide 6, derived from the cell-cell adhesion protein encoded by Cdh16), and one without functional annotation (peptide 10, derived from the kelch-domain containing protein encoded by Klhdc7a) (see Supplemental Table 3). Peptides 1 and 7 are derived from cytoplasmic regions of their respective proteins, while peptide 6 is derived from the extracellular region. These proteins may be adducted by IsoLG and processed via several pathways. Renal epithelial cells experience increased oxidative stress in hypertension (51, 52), and thus these proteins could undergo IsoLG adduction in their cell of origin. It is also possible that previously unadducted proteins, released with cellular debris or upon cell death in the kidney, are engulfed by DCs where they are then IsoLG-adducted. In keeping with this, we have shown that superoxide production is increased by more than 6-fold in DCs of hypertensive mice and this is absent in mice lacking NOX2 (18).

IsoLGs are a family of 8 regioisomers, and reaction of IsoLGs with lysine form a number of adduct species including pyrrole and oxidized pyrroles (53). Chromatograms derived from mass-spectrometry data obtained one of the immunogenic IsoLG-adducted peptides revealed characteristic signals congruent with different IsoLG adducts including anhydrolactams and andihydropyrroles (Supplemental Figure 8). While such differences in IsoLG structure may not appreciably change their affinity for MHC-I, different IsoLG regioisomer or adduct species could select for different T cell populations. This mechanism might explain why we observe many distinct T cell clonotypes in the kidneys of hypertensive mice, rather an a few single dominant clonal populations (9). Additionally, the robust T cell response observed despite a relatively low amount of IsoLG-adducted peptide (as assessed by mass spectrometry, Supplemental Figure 7) suggests that the adducts may be especially immunogenic.

Our data strongly suggests that IsoLG-adducts are restricted to certain MHC-I variants in mice and preferentially displayed by certain HLA variants in humans. Such data implies there might exist a correlation between “high risk” HLA variants and hypertension as is the case for other diseases (54). There are several studies with small populations that correlate specific class-II HLA alleles with hypertension severity, though this may be due to linkage disequilibrium rather than HLA itself (55, 56). A more recent study correlating phenotypes with imputed HLA alleles in a large clinical database do not reproduce these associations, and there is no single class-I HLA that confers significantly elevated risk for hypertension (57). This may be due in part to the “mosaic” nature of hypertension, where distinct pathogenic stimuli act in concert to mediate a common phenotype (58). While single HLA alleles may be insufficient to promote disease it is also possible that combinations of multiple alleles, or haplotypes, may confer either risk (if all are predisposed to IsoLG-adduct presentation) or protection (if they have low affinity for IsoLG adducts). The computational modeling pipeline employed for this study could also be employed to screen large numbers of antigen/HLA combinations. Further studies combining this screening tool with large databases containing both imputed HLA alleles and diagnosis codes may help identify high risk haplotypes, and individual alleles later screened for IsoLG-adduct affinity *in vitro*.

There are several important limitations to consider for this work. Firstly, the peptides we screened are N-terminal acetylated. While necessary to prevent IsoLG adduction at the peptide N-terminus, N-terminal acetylation can also alter peptide binding affinity for MHC-I resulting in false negatives in our screening studies (59). Solid-phase peptide synthesis using pre-adducted lysine residues could circumvent this problem, but an optimized protocol for manufacturing such residues is not yet available. In our initial computational screen, we also excluded peptides predicted to have a low binding affinity for H-2D^b^. This exclusion step reduced the number of peptides included in the screen by several orders of magnitude, but also may have rejected peptides with increased binding affinity for MHC-I following IsoLG-adduction.

While we exclusively considered CD8^+^ T cell mediated responses to peptide candidates, it is also possible that CD4^+^ T cells recognize these same or similar antigens presented by MHC-II. IL-17 producing CD4^+^ T cells and gamma delta T cells are an important source of inflammation in hypertension and recognize IsoLG-adducted peptides in mouse models of nonischemic heart failure (8, 17, 60). Future studies should examine the ability of these cells to recognize similar IsoLG-adducted peptides.

Despite these limitations, identifying antigens that mediate T cell inflammation in hypertension is an important discovery with broad implications. Future work expanding the scope of immunogenic IsoLG-adducted self-peptides, identifying the T cells that recognize them, and characterizing molecularly similar antigens from human pathogens that may evoke cross-reactive memory T cell responses will greatly enhance our understanding of how inflammation drives morbidity and mortality in this increasingly common ailment.

## Methods

### Transgenic animals and murine hypertension models

DNA fragments containing the H-2K^b^ and H-2D^b^ heavy chains with 6-His tags and without transmembrane domain regions downstream of a CD11c promoter were amplified and cloned into the pcDNA3.1 (H-2D^b^) and pET-3a (H-2K^b^) vectors. Resultant vectors were transformed into ampicillin-treated *E. Coli* (DH5α), purified using the ThermoFisher PureLink Maxiprep kit, and sequenced to confirmed insertion. The resultant constructs were used to perform pronuclear injections for creation of transgenic C57Bl/6 mice with soluble forms of H2-D^b^ and H2-K^b^ respectively.

To induce hypertension, mice were implanted with osmotic minipumps (Alzet) containing angiotensin II delivered at a rate of 490 ng/min/kg (regular dosing) or 140 ng/min/kg (subpressor dosing), or sodium acetate buffer (sham controls). Mice were studied at 12 weeks of age and received angiotensin II for two weeks, and blood pressure tail cuff measurements using the MC4000 Multichannel System for mice (Hatteras, Inc). In some experiments, 2-HOBA was provided concurrently in drinking water at a concentration of 1g/L. Animals were euthanized by CO_2_ asphyxiation prior to tissue isolation. All animal experiments were approved by the Vanderbilt University Medical Center Institutional Animal Care and Use Committee.

### In silico studies and computational screening

Rosetta stable release version 3.13 was used for all computational studies with FlexPepDock refinement. We created Rosetta params files for modeling isolevuglandin-adducted lysine as previously described (61). Preliminary structures were generated by identifying the best-scoring pre-existing template structure available in the PDB (after alignment and scoring with the Blocks Substitution Matrix BLOSUM62) and mutating peptide residue to match those of the query sequencing using the Rosetta Scripts mover MutateResidue prior to prepacking (62). MHC-I models were generated using AlphaFold2 (63). Unless otherwise specified, 250 models were generated for each peptide/MHC-I complex using computational resources available through the Vanderbilt Advanced Computing Center for Research and Education (ACCRE). The Rosetta energy composite score reweighted_sc, which provides additional weighting to interactions at the peptide-receptor interface, was averaged from the top 5 scoring models. PyMOL molecular graphics software was used to visualize the generated models (Schrödinger). All code used to generate the models in this manuscript is provided in a public GitHub repository. Candidate peptides nine residues in length were derived from the sequences of proteins overexpressed in renal tissue. We queried the Human Protein Atlas for proteins with a greater than 4-fold increase in RNA expression in the kidney compared to other tissues, identified mouse homologs, and used NetMHCpan 4.0 to identify peptides derived from those sequences that were likely to bind strongly to H-2D^b^ (40).

### Peptide synthesis, IsoLG adduction, and in vitro IsoLG adduct generation

Candidate peptides commercially produced (EZBiolab). Peptides were N-terminal acetylated to prevent unwanted N-terminus IsoLG adduction, and purity confirmed with high performance liquid chromatography. IsoLG was synthesized as described (64). Peptides were incubated with IsoLG at a 1:1 molar ratio of lysine to IsoLG overnight at 4°C in aqueous solution to generate adducts as previously described (14). Unadducted and IsoLG-adducted peptides were stored at 4°C at a concentration of 1mM until used in cell culture or flow cytometry.

Native IsoLG generation was induced by treating cultured cells with tert-butyl hydroperoxide (tBHP, Sigma-Adrich) at 100 μM for 30 minutes as previously described (18). After treatment, cells were centrifuged at 350 g for 5 minutes and the media replaced. Cells were incubated overnight prior to further analysis.

### Cell isolation and culture

For assays with bead-bound MHC-I, spleens from H-2D^b^ or H-2K^b^ transgenic mice were isolated, passed through a 70 μm mesh, and red blood cells lysed (RBC lysis buffer, eBioscience) prior to culture in RPMI supplemented with 10% v/v fetal bovine serum, 1% v/v penicillin/streptomycin (Gibco), and 2 μL/500mL of β-mercaptoethanol (Sigma-Aldrich). After 72 hours, media was collected and incubated with nickel agarose beads (ThermoFisher) and the beads collected through centrifugation before being added to T cell cultures. If indicated, a single-chain variable fragment antibody that recognizes IsoLG-adducts (D11) was added to cultures simultaneously (65).

For T cell proliferation assays, dendritic cells were isolated from the spleens of C57BL/6 animals using the Miltenyi Pan Dendritic Cell Isolation Kit and cultured in RPMI supplemented with 10% v/v fetal bovine serum, 10 mM HEPES buffer, 1% v/v sodium pyruvate (Gibco), 1% v/v penicillin/streptomycin (Gibco), and 2 μL/500mL of β-mercaptoethanol (Sigma-Aldrich). Adducted or unadducted peptides were added to DC cultures overnight at a concentration of 10 μM, DCs centrifuged, and supernatant removed prior to the addition of T cells. CD3^+^ cells were isolated from the bone marrow of adult C57BL/6 angiotensin II or sham treated mice using the Miltenyi Pan T Cell Isolation Kit II and cultured in RPMI supplemented with 10% v/v fetal bovine serum, 1% v/v MEM non-essential amino acids solution (Gibco), 1% v/v penicillin/streptomycin (Gibco), and 0.4 uL/100mL of β-mercaptoethanol (Sigma-Aldrich). T cells were stained with CellTrace CFSE (ThermoFisher) prior to incubation with DCs to track proliferation. A 1:1000 dilution of anti-CD28 was added to T cell and DC co-cultures for co-stimulation. T cells were cultured for five days prior to analysis with flow cytometry as described.

### Tissue isolation and flow cytometry

Femurs and tibias, aortas, and kidneys were isolated from sham or angiotensin II treated mice following perfusion of a normal saline solution to remove residual leukocytes from the peripheral blood. Long bones were flushed with phosphate buffered saline to isolate bone marrow cells. Aortas were manually homogenized and digested for 20 minutes at 37°C in RPMI with 1 mg/mL of collagenase A and collagenase B and 0.1 mg/mL DNAse I (Roche). Kidneys were homogenized using the Miltenyi gentleMACS Dissociator and digested for 20 minutes at 37°C in RPMI with 2 mg/mL collagenase D (Roche) and 0.1 mg/mL DNAse I. Red blood cells were lysed for all tissues using RBC lysis buffer (eBioscience). Single cell suspensions were stained with Live/Dead violent fluorescent reactive dye (ThermoFisher), Fc receptor block, and stained with antibodies for CD3 (BV650), CD4 (FITC), CD8 (APC), CD44 (APC-Fire750), and CD62L (PE-Fire810) (BioLegend). To identify peptide-specific T cells, suspensions were also stained with H-2D^b^/IgG1 fusion protein loaded with unadducted or IsoLG-adducted peptide (160:1 molar ratio overnight at 37°C) at 1 ug per sample. Secondary staining to identify H-2D^b^/IgG1-bound cells was done with anti-mouse IgG1 (PE) (BD Biosciences). Samples were run on a Cytek Aurora 4-laser flow cytometer and analyzed using FlowJo. A minimum of 1×10^6^ events were collected and cell counts determined based on total cell number (bone marrow and aorta cells, determined after isolation using a hemocytometer with trypan blue staining) or normalized to organ weight (kidney).

### Assessment of IsoLG adduct presentation by single-variant HLA-expressing cells

We identified HLA alleles to screen for IsoLG adduct presentation by selecting the ten HLA alleles with the highest phenotypic frequency present in three populations of the US National Merit Donor Program available through the Allele Frequency Net Database to best capture haplotype diversity (42). The populations included USA NMDP European Caucasian, USA NMDP Chinese, and USA NMDP African American Pop 2. Twenty-two non-duplicate HLA-A, -B, and -C alleles were identified in this manner.

The HLA-null human cell line K562 was transduced with each HLA allele with either an adenovirus vector or electroporation in conjunction with a transposon sequence and sleeping beauty transposase. After transduction and expansion, HLA-expressing cells were enriched with a positive magnetic sort using the biotinylated pan-HLA antibody (clone W6/32, Biolegend) in conjunction with the ThermoFisher CELLection Dynabead cell isolation kit. HLA expression was confirmed with flow cytometry after staining with Live/Dead violent fluorescent reactive dye and anti-pan-HLA (BV650). Non-transduced K562 cells were used as a null control.

To induce IsoLG adduct presentation, cells were treated with tBHP as described. If indicated, cells were also maintained in media containing ethyl-2-HOBA at a concentration of 200 μM for the duration of the experiment. After incubation overnight, cells were stained with live/dead violet fluorescent reactive dye and anti-pan-HLA (BV650) and biotinylated D11 conjugated to streptavidin APC-Cy7. D11 was biotinylated prior to conjugation with the streptavidin fluorophore using the Lightning Link A Biotinylation Kit (Abcam).

## Acknowledgements

This work was conducted in part using the resources of the Advanced Computing Center for Research and Education at Vanderbilt University, Nashville, TN. Mice were created in the Vanderbilt Genome Editing Resource Center.

