## Supplemental Materials for "Discovery of post-translationally modified self-peptides that promote hypertension"

**H-2D^b^ (912 bp)**

atgggggcgatggctccgcgcacgctgctcctgctgctggcggccgccctggccccgactcagacccgcgcgggcccacactcgatgcggtatttcgagaccgccgtgtcccggcccggcctcgaggagccccggtacatctctgtcggctatgtggacaacaaggagttcgtgcgcttcgacagcgacgcggagaatccgagatatgagccgcgggcgccgtggatggagcaggaggggccggagtattgggagcgggaaacacagaaagccaagggccaagagcagtggttccgagtgagcctgaggaacctgctcggctactacaaccagagcgcgggcggctctcacacactccagcagatgtctggctgtgacttggggtcggactggcgcctcctccgcgggtacctgcagttcgcctatgaaggccgcgattacatcgccctgaacgaagacctgaaaacgtggacggcggcggacatggcggcgcagatcacccgacgcaagtgggagcagagtggtgctgcagagcattacaaggcctacctggagggcgagtgcgtggagtggctccacagatacctgaagaacgggaacgcgacgctgctgcgcacagattccccaaaggcacatgtgacccatcaccccagatctaaaggtgaagtcaccctgaggtgctgggccctgggcttctaccctgctgacatcaccctgacctggcagttgaatggggaggagctgacccaggacatggagcttgtggagaccaggcctgcaggggatggaaccttccagaagtgggcatctgtggtggtgcctcttgggaaggagcagaattacacatgccgtgtgtaccatgaggggctgcctgagcccctcaccctgagatgggagcctcctccgtccact

**H-2K^b^ (840 bp)**

ggcccacactcgctgaggtatttcgtcaccgccgtgtcccggcccggcctcggggagccccggtacatggaagtcggctacgtggacgacacggagttcgtgcgcttcgacagcgacgcggagaatccgagatatgagccgcgggcgcggtggatggagcaggaggggcccgagtattgggagcgggagacacagaaagccaagggcaatgagcagagtttccgagtggacctgaggaccctgctcggctactacaaccagagcaagggcggctctcacactattcaggtgatctctggctgtgaagtggggtccgacgggcgactcctccgcgggtaccagcagtacgcctacgacggctgcgattacatcgccctgaacgaagacctgaaaacgtggacggcggcggacatggcggcgctgatcaccaaacacaagtgggagcaggctggtgaagcagagagactcagggcctacctggagggcacgtgcgtggagtggctccgcagatacctgaagaacgggaacgcgacgctgctgcgcacagattccccaaaggcccatgtgacccatcacagcagacctgaagataaagtcaccctgaggtgctgggccctgggcttctaccctgctgacatcaccctgacctggcagttgaatggggaggagctgatccaggacatggagcttgtggagaccaggcctgcaggggatggaaccttccagaagtgggcatctgtggtggtgcctcttgggaaggagcagtattacacatgccatgtgtaccatcaggggctgcctgagcccctcaccctgagatgggagcctcctccatccact

Supplemental Table 1: Soluble MHC-I heavy chain sequences for H-2D^b^ and H-2K^b^

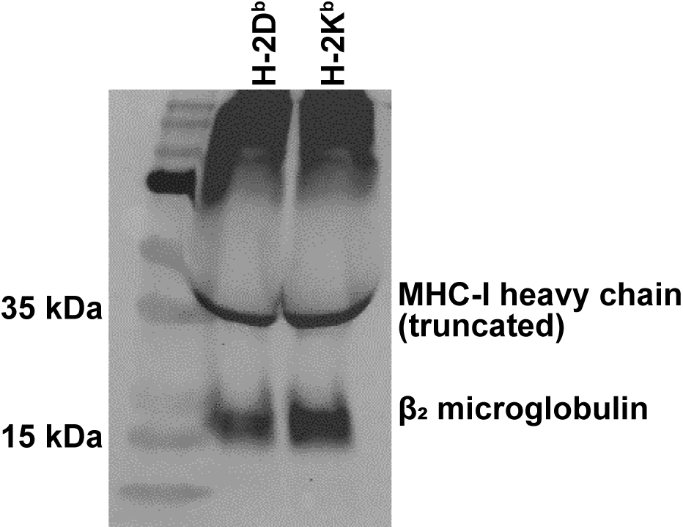

Supplemental Figure 1: Soluble His-tagged MHC-I is present in the media of cultured splenocytes isolated from transgenic mice. His-tagged proteins were immunoprecipitated and separated by gel electrophoresis. Staining with antibodies for MHC-I and β2 microglobulin confirms the presence of soluble MHC-I.

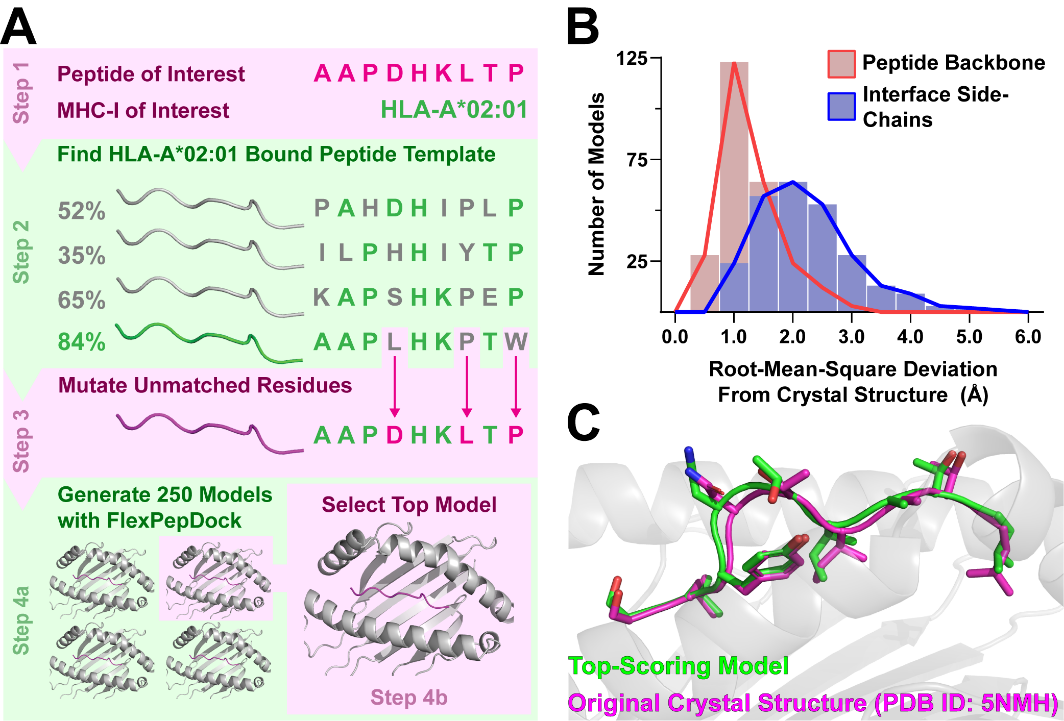

Supplemental Figure 2: FlexPepDock refinement can accurately recapitulate MHC-I bound peptide structures using pre-existing templates. (A) Diagram illustrating our approach for selecting peptide templates from the PDB based on sequence similarity. (B) A benchmark on available MHC-I molecules in the PDB with a resolution <2.5 Å yields structures with low root-mean-square deviation (RMSD) from the original crystal structures (n=329). Heavy atom side-chain RMSD was also <2.5 for most structures generated. (C) Example of a peptide model generated by FlexPepDock refinement (PDB ID 5NMH. Model shown in green, original structure in magenta).

|  |  |  | **Template** | |  | **RMSD** | |
| --- | --- | --- | --- | --- | --- | --- | --- |
| **PDB** | **MHC-I** | **Epitope** | **PDB** | **MHC-I** |  | **Backbone** | **Interface** |
| 1A1M | B*53:01 | TPYDINQML | 1A1O | B*53:01 |  | 1.35 | 2.59 |
| 1A1N | B*35:01 | VPLRPMTY | 3BWA | B*35:08 |  | 1.73 | 2.48 |
| 1A1O | B*53:01 | KPIVQYDNF | 1A1M | B*53:01 |  | 1.08 | 2.94 |
| 1A9E | B*35:01 | LPPLDITPY | 4QRR | B*35:01 |  | 1.98 | 3.24 |
| 1AGB | B*08:01 | GGRKKYKL | 1AGD | B*08:01 |  | 0.79 | 1.18 |
| 1AGC | B*08:01 | GGKKKYQL | 1AGD | B*08:01 |  | 0.40 | 1.33 |
| 1AGD | B*08:01 | GGKKKYKL | 1AGE | B*08:01 |  | 0.41 | 1.55 |
| 1AGE | B*08:01 | GGKKKYRL | 1AGD | B*08:01 |  | 0.36 | 1.47 |
| 1AGF | B*08:01 | GGKKRYKL | 1AGD | B*08:01 |  | 0.45 | 1.43 |
| 1DUY | A*02:01 | LFGYPVYV | 4F7T | A*24:02 |  | 3.06 | 4.63 |
| 1DUZ | A*02:01 | LLFGYPVYV | 3QFJ | A*02:01 |  | 1.47 | 1.87 |
| 1E27 | B*51:01 | LPPVVAKEI | 1A1M | B*53:01 |  | 2.77 | 4.14 |
| 1EEY | A*02:01 | ILSALVGIV | 1EEZ | A*02:01 |  | 1.37 | 1.95 |
| 1EEZ | A*02:01 | ILSALVGIL | 1EEY | A*02:01 |  | 1.32 | 1.83 |
| 1EFX | C*03:04 | GAVDPLLAL | 4NT6 | C*08:01 |  | 2.27 | 3.59 |
| 1HHG | A*02:01 | TLTSCNTSV | 7N1E | A*02:01 |  | 1.60 | 2.22 |
| 1HHH | A*02:01 | FLPSDFFPSV | 5C0G | A*02:01 |  | 2.21 | 3.43 |
| 1HSA | B*27:05 | ARAAAAAAA | 1JGE | B*27:05 |  | 1.52 | 1.67 |
| 1I1F | A*02:01 | FLKEPVHGV | 1I1Y | A*02:01 |  | 0.90 | 1.90 |
| 1I1Y | A*02:01 | YLKEPVHGV | 1I1F | A*02:01 |  | 1.17 | 2.58 |
| 1I4F | A*02:01 | GVYDGREHTV | 6TRN | A*02:01 |  | 0.82 | 2.63 |
| 1I7R | A*02:01 | FAPGFFPYL | 1LP9 | A*02:01 |  | 1.13 | 1.72 |
| 1I7T | A*02:01 | ALWGVFPVL | 1LP9 | A*02:01 |  | 0.79 | 1.01 |
| 1I7U | A*02:01 | ALWGFVPVL | 1LP9 | A*02:01 |  | 1.38 | 2.12 |
| 1JF1 | A*02:01 | ELAGIGILTV | 4JFE | A*02:01 |  | 0.82 | 1.55 |
| 1JGE | B*27:05 | GRFAAAIAK | 1HSA | B*27:05 |  | 0.77 | 1.64 |
| 1LP9 | A*02:01 | ALWGFFPVL | 1I7U | A*02:01 |  | 1.00 | 1.45 |
| 1M05 | B*08:01 | FLRGRAYGL | 3SKM | B*08:01 |  | 1.84 | 2.06 |
| 1M6O | B*44:02 | EEFGRAFSF | 3L3G | B*44:02 |  | 0.59 | 1.34 |
| 1N2R | B*44:03 | EEFGRAFSF | 3KPN | B*44:03 |  | 1.46 | 3.32 |
| 1OGA | A*02:01 | GILGFVFTL | 5HHQ | A*02:01 |  | 1.56 | 2.30 |
| 1Q94 | A*11:01 | AIFQSSMTK | 6JOZ | A*11:01 |  | 1.50 | 2.23 |
| 1QQD | C*04:01 | QYDDAVYKL | 5VGD | C*05:01 |  | 1.16 | 2.31 |
| 1QRN | A*02:01 | LLFGYAVYV | 1DUZ | A*02:01 |  | 1.09 | 2.95 |
| 1QSE | A*02:01 | LLFGYPRYV | 1DUZ | A*02:01 |  | 1.20 | 1.57 |
| 1QSF | A*02:01 | LLFGYPVAV | 1DUZ | A*02:01 |  | 1.06 | 2.61 |
| 1QVO | A*11:01 | QVPLRPMTYK | 5WKF | A*11:01 |  | 1.98 | 3.23 |
| 1S8D | A*02:01 | SLANTVATL | 2V2X | A*02:01 |  | 1.45 | 2.59 |
| 1S9X | A*02:01 | SLLMWITQA | 1S9Y | A*02:01 |  | 1.39 | 2.29 |
| 1S9Y | A*02:01 | SLLMWITQS | 1S9X | A*02:01 |  | 0.99 | 1.80 |
| 1SYS | B*44:03 | EEPTVIKKY | 3KPN | B*44:03 |  | 1.94 | 2.16 |
| 1SYV | B*44:05 | EEFGRAFSF | 3KPP | B*44:05 |  | 1.65 | 3.34 |
| 1T1X | A*02:01 | SLYLTVATL | 2V2W | A*02:01 |  | 0.98 | 1.75 |
| 1T1Y | A*02:01 | SLYNVVATL | 2V2W | A*02:01 |  | 0.96 | 1.40 |
| 1T1Z | A*02:01 | ALYNTAAAL | 2V2W | A*02:01 |  | 0.81 | 1.29 |
| 1TVB | A*02:01 | ITDQVPFSV | 1TVH | A*02:01 |  | 0.71 | 2.03 |
| 1TVH | A*02:01 | IMDQVPFSV | 6VMC | A*02:01 |  | 0.79 | 2.54 |
| 1UXS | B*27:05 | RRRWRRLTV | 5IB2 | B*27:05 |  | 1.38 | 2.53 |
| 1W0V | B*27:05 | RRLPIFSRL | 2BSR | B*27:05 |  | 1.09 | 1.93 |
| 1X7Q | A*11:01 | KTFPPTEPK | 1Q94 | A*11:01 |  | 2.04 | 3.27 |
| 1XR8 | B*15:01 | LEKARGSTY | 1XR9 | B*15:01 |  | 0.96 | 2.97 |
| 1XR9 | B*15:01 | ILGPPGSVY | 1XR8 | B*15:01 |  | 1.76 | 2.26 |
| 2A83 | B*27:05 | RRRWHRWRL | 5IB2 | B*27:05 |  | 1.01 | 2.47 |
| 2AXF | B*35:08 | APQPAPENAY | 5VZ5 | B*15:01 |  | 2.77 | 3.65 |
| 2AXG | B*35:01 | APQPAPENAY | 5VZ5 | B*15:01 |  | 2.54 | 3.46 |
| 2BCK | A*24:02 | VYGFVRACL | 6XQA | A*24:02 |  | 1.07 | 1.67 |
| 2BNQ | A*02:01 | SLLMWITQV | 3KLA | A*02:01 |  | 0.91 | 2.63 |
| 2BSR | B*27:05 | RRIYDLIEL | 1W0V | B*27:05 |  | 1.26 | 2.38 |
| 2BST | B*27:05 | SRYWAIRTR | 3LV3 | B*27:05 |  | 1.02 | 3.09 |
| 2BVP | B*57:03 | ISPRTLNAW | 5VVP | B*57:03 |  | 1.44 | 2.25 |
| 2BVQ | B*57:03 | KAFSPEVI | 3W39 | B*52:01 |  | 0.98 | 1.60 |
| 2CIK | B*35:01 | KPIVVLHGY | 4QRR | B*35:01 |  | 1.43 | 2.68 |
| 2CLR | A*02:01 | MLLSVPLLLG | 4JFP | A*02:01 |  | 2.82 | 3.92 |
| 2GT9 | A*02:01 | EAAGIGILTV | 1JF1 | A*02:01 |  | 1.03 | 1.48 |
| 2GTW | A*02:01 | LAGIGILTV | 3QFD | A*02:01 |  | 2.86 | 3.31 |
| 2GTZ | A*02:01 | ALGIGILTV | 3QFD | A*02:01 |  | 1.22 | 1.83 |
| 2HN7 | A*11:01 | AIMPARFYPK | 5WKF | A*11:01 |  | 2.09 | 3.87 |
| 2P5E | A*02:01 | SLLMWITQC | 1S9X | A*02:01 |  | 0.81 | 2.05 |
| 2V2W | A*02:01 | SLYNTVATL | 5NMH | A*02:01 |  | 0.87 | 1.49 |
|  |  |  | **Template** | |  | **RMSD** | |
| **PDB** | **MHC-I** | **Epitope** | **PDB** | **MHC-I** |  | **Backbone** | **Interface** |
| 2X4O | A*02:01 | KLTPLCVTL | 3MRC | A*02:01 |  | 1.58 | 2.60 |
| 2X4S | A*02:01 | AMDSNTLEL | 3D25 | A*02:01 |  | 1.25 | 2.40 |
| 2X4U | A*02:01 | ILKEPVHGV | 1I1F | A*02:01 |  | 0.94 | 1.49 |
| 2XPG | A*03:01 | KLIETYFSK | 3RL1 | A*03:01 |  | 1.05 | 2.50 |
| 3BO8 | A*01:01 | EADPTGHSY | 5BRZ | A*01:01 |  | 0.75 | 1.33 |
| 3BP4 | B*27:05 | IRAAPPPLF | 1HSA | B*27:05 |  | 1.75 | 2.74 |
| 3BWA | B*35:08 | FPTKDVAL | 1A1N | B*35:01 |  | 0.81 | 1.47 |
| 3BXN | B*14:02 | IRAAPPPLF | 4O2E | B*39:01 |  | 2.43 | 3.26 |
| 3C9N | B*15:01 | VQQESSFVM | 1XR8 | B*15:01 |  | 1.07 | 1.97 |
| 3D25 | A*02:01 | VLHDDLLEA | 3FT4 | A*02:01 |  | 0.86 | 1.66 |
| 3DX6 | B*44:02 | EENLLDFVRF | 2AXF | B*35:08 |  | 1.42 | 2.67 |
| 3DX7 | B*44:03 | EENLLDFVRF | 2AXG | B*35:01 |  | 2.02 | 4.28 |
| 3DX8 | B*44:05 | EENLLDFVRF | 2AXF | B*35:08 |  | 2.25 | 4.54 |
| 3FQT | A*02:01 | GLLGSPVRA | 1QSF | A*02:01 |  | 1.50 | 2.92 |
| 3FT4 | A*02:01 | VLRDDLLEA | 3D25 | A*02:01 |  | 0.76 | 1.60 |
| 3GIV | A*02:01 | SLFNTVATLY | 3MRN | A*02:01 |  | 2.23 | 5.27 |
| 3GSQ | A*02:01 | NLVPSVATV | 3GSU | A*02:01 |  | 0.84 | 1.32 |
| 3GSR | A*02:01 | NLVPVVATV | 6Q3K | A*02:01 |  | 0.82 | 1.61 |
| 3GSU | A*02:01 | NLVPTVATV | 3GSQ | A*02:01 |  | 0.82 | 1.22 |
| 3GSV | A*02:01 | NLVPQVATV | 3GSQ | A*02:01 |  | 0.86 | 1.68 |
| 3GSW | A*02:01 | NLVPMVAAV | 3GSX | A*02:01 |  | 0.83 | 1.85 |
| 3GSX | A*02:01 | NLVPMVAVV | 3GSW | A*02:01 |  | 0.86 | 1.58 |
| 3H7B | A*02:01 | MLWGYLQYV | 1QSE | A*02:01 |  | 0.90 | 1.45 |
| 3HPJ | A*02:01 | RMFPNAPYL | 1QRN | A*02:01 |  | 1.05 | 2.87 |
| 3I6G | A*02:01 | GLMWLSYFV | 5HHN | A*02:01 |  | 0.99 | 1.85 |
| 3I6K | A*02:01 | TLACFVLAAV | 4JFQ | A*02:01 |  | 1.89 | 2.71 |
| 3KLA | A*02:01 | SLLMWITQL | 2BNQ | A*02:01 |  | 1.22 | 2.29 |
| 3KPL | B*44:02 | EEYLQAFTY | 3KPM | B*44:02 |  | 1.40 | 2.75 |
| 3KPM | B*44:02 | EEYLKAWTF | 3KPL | B*44:02 |  | 1.28 | 3.12 |
| 3KPN | B*44:03 | EEYLQAFTY | 3KPO | B*44:03 |  | 1.69 | 2.88 |
| 3KPO | B*44:03 | EEYLKAWTF | 3KPN | B*44:03 |  | 1.71 | 3.62 |
| 3KPP | B*44:05 | EEYLQAFTY | 3KPQ | B*44:05 |  | 1.33 | 2.62 |
| 3KPQ | B*44:05 | EEYLKAWTF | 3KPP | B*44:05 |  | 1.45 | 3.15 |
| 3L3D | B*44:02 | EEAGRAFSF | 1M6O | B*44:02 |  | 0.66 | 2.00 |
| 3L3G | B*44:02 | EEFGAAFSF | 1M6O | B*44:02 |  | 0.87 | 1.81 |
| 3L3I | B*44:02 | EEFGRAASF | 1M6O | B*44:02 |  | 0.87 | 2.33 |
| 3L3J | B*44:02 | EEAGAAFSF | 3L3D | B*44:02 |  | 1.11 | 1.69 |
| 3L3K | B*44:02 | EEFGAAASF | 3L3I | B*44:02 |  | 1.50 | 1.63 |
| 3LKN | B*35:01 | LPFERATIM | 3LKR | B*35:01 |  | 0.95 | 1.60 |
| 3LKO | B*35:01 | LPFDRTTIM | 3LKQ | B*35:01 |  | 0.68 | 2.03 |
| 3LKP | B*35:01 | LPFDKSTIM | 3LKQ | B*35:01 |  | 0.90 | 1.87 |
| 3LKQ | B*35:01 | LPFDKTTIM | 3LKO | B*35:01 |  | 0.73 | 1.06 |
| 3LKR | B*35:01 | LPFERATVM | 3LKN | B*35:01 |  | 0.73 | 1.28 |
| 3LKS | B*35:01 | LPFEKSTVM | 3LKP | B*35:01 |  | 0.86 | 1.32 |
| 3LV3 | B*27:05 | SRRWRRWNR | 5IB2 | B*27:05 |  | 1.44 | 4.15 |
| 3MGO | A*02:01 | RLYQNPTTYI | 3MGT | A*02:01 |  | 1.65 | 2.96 |
| 3MGT | A*02:01 | KLYQNPTTYI | 3MGO | A*02:01 |  | 1.99 | 3.08 |
| 3MR9 | A*02:01 | NLVPAVATV | 3GSQ | A*02:01 |  | 0.82 | 1.30 |
| 3MRB | A*02:01 | NLVPMVHTV | 6Q3K | A*02:01 |  | 0.95 | 1.38 |
| 3MRC | A*02:01 | NLVPMCATV | 6Q3K | A*02:01 |  | 1.93 | 2.72 |
| 3MRD | A*02:01 | NLVPMGATV | 3MRC | A*02:01 |  | 1.34 | 1.70 |
| 3MRE | A*02:01 | GLCTLVAML | 3GSX | A*02:01 |  | 1.60 | 2.06 |
| 3MRG | A*02:01 | CINGVCWTV | 3MRJ | A*02:01 |  | 1.54 | 2.25 |
| 3MRH | A*02:01 | CISGVCWTV | 3MRG | A*02:01 |  | 0.91 | 2.24 |
| 3MRI | A*02:01 | CINMWCWTV | 3MRJ | A*02:01 |  | 1.43 | 3.73 |
| 3MRJ | A*02:01 | CINGMCWTV | 3MRG | A*02:01 |  | 1.02 | 1.88 |
| 3MRK | A*02:01 | PLFQVPEPV | 6VMC | A*02:01 |  | 1.24 | 2.54 |
| 3MRL | A*02:01 | CINGVVWTV | 3MRG | A*02:01 |  | 0.99 | 2.26 |
| 3MRM | A*02:01 | KLVALGINAV | 3MRP | A*02:01 |  | 1.18 | 1.79 |
| 3MRN | A*02:01 | LLFNILGGWV | 3GIV | A*02:01 |  | 2.76 | 4.91 |
| 3MRO | A*02:01 | ELAGWGILTV | 1JF1 | A*02:01 |  | 0.98 | 2.28 |
| 3MRP | A*02:01 | ELAGLGINTV | 4JFQ | A*02:01 |  | 0.94 | 1.62 |
| 3MRR | A*02:01 | LLAGIGTVPI | 6AM5 | A*02:01 |  | 1.26 | 1.77 |
| 3OX8 | A*02:03 | FLPSDFFPSV | 5C0G | A*02:01 |  | 1.61 | 3.36 |
| 3OXR | A*02:06 | FLPSDFFPSV | 5C0G | A*02:01 |  | 2.12 | 4.13 |
| 3OXS | A*02:07 | FLPSDFFPSV | 5C0G | A*02:01 |  | 1.93 | 3.68 |
| 3PWJ | A*02:01 | LLYGFVNYV | 3PWN | A*02:01 |  | 1.05 | 1.49 |
| 3PWL | A*02:01 | LGYGFVNYI | 3PWN | A*02:01 |  | 1.01 | 2.15 |
| 3PWN | A*02:01 | LLYGFVNYI | 3PWJ | A*02:01 |  | 0.89 | 1.33 |
|  |  |  | **Template** | |  | **RMSD** | |
| **PDB** | **MHC-I** | **Epitope** | **PDB** | **MHC-I** |  | **Backbone** | **Interface** |
| 3QFJ | A*02:01 | LLFGFPVYV | 1DUZ | A*02:01 |  | 0.91 | 2.47 |
| 3REW | A*02:01 | CLGGLLTMV | 3MRL | A*02:01 |  | 1.57 | 2.28 |
| 3RL1 | A*03:01 | AIFQSSMTK | 7L1C | A*03:01 |  | 1.11 | 2.34 |
| 3RL2 | A*03:01 | QVPLRPMTYK | 5WKF | A*11:01 |  | 2.77 | 4.00 |
| 3SKM | B*08:01 | FLRGRAYVL | 1M05 | B*08:01 |  | 1.89 | 3.08 |
| 3SPV | B*08:01 | RAKFKQLL | 1AGD | B*08:01 |  | 0.53 | 1.07 |
| 3UPR | B*57:01 | HSITYLLPV | 5U98 | B*57:01 |  | 1.20 | 2.22 |
| 3UTQ | A*02:01 | ALWGPDPAAA | 5C0D | A*02:01 |  | 1.16 | 1.46 |
| 3VH8 | B*57:01 | LSSPVTKSF | 5VUD | B*57:01 |  | 1.14 | 1.24 |
| 3VRJ | B*57:01 | LTTKLTNTNI | 5T6X | B*57:01 |  | 3.42 | 5.10 |
| 3VXN | A*24:02 | RYPLTFGWCF | 5HGD | A*24:02 |  | 1.52 | 2.23 |
| 3VXS | A*24:02 | RYPLTLGWCF | 3VXN | A*24:02 |  | 1.06 | 1.60 |
| 3W39 | B*52:01 | TAFTIPSI | 2BVQ | B*57:03 |  | 1.24 | 2.13 |
| 3WL9 | A*24:02 | NYTPGPGIRF | 3WLB | A*24:02 |  | 0.90 | 1.41 |
| 3WLB | A*24:02 | NYTPGPGTRF | 3WL9 | A*24:02 |  | 1.99 | 1.59 |
| 4F7M | A*24:02 | LYASPQLEGF | 7JYU | A*24:02 |  | 1.06 | 2.05 |
| 4F7T | A*24:02 | RYGFVANF | 5HGB | A*24:02 |  | 0.98 | 1.75 |
| 4G8I | B*27:05 | KRWIIMGLNK | 4G9D | B*27:05 |  | 1.29 | 1.31 |
| 4G9D | B*27:05 | KRWIILGLNK | 4G8I | B*27:05 |  | 2.89 | 4.09 |
| 4GKN | A*02:01 | FATGIGIITV | 4GKS | A*02:01 |  | 1.65 | 2.54 |
| 4GKS | A*02:01 | FLTGIGIITV | 4GKN | A*02:01 |  | 1.25 | 2.16 |
| 4HWZ | A*68:01 | AIFQSSMTK | 4HX1 | A*68:02 |  | 1.56 | 2.48 |
| 4HX1 | A*68:02 | SVYDFFVWL | 4I48 | A*68:02 |  | 1.30 | 2.57 |
| 4I48 | A*68:02 | TLTSCNTSV | 4HX1 | A*68:02 |  | 1.90 | 2.37 |
| 4I4W | A*02:01 | ILAKFLHWL | 5MEP | A*02:01 |  | 1.10 | 1.77 |
| 4JFE | A*02:01 | ELAGIGALTV | 1JF1 | A*02:01 |  | 0.93 | 1.48 |
| 4JFO | A*02:01 | ALAGIGILTV | 1JF1 | A*02:01 |  | 0.82 | 1.70 |
| 4JFP | A*02:01 | ELAAIGILTV | 1JF1 | A*02:01 |  | 0.99 | 1.51 |
| 4JFQ | A*02:01 | ELAGIGIATV | 1JF1 | A*02:01 |  | 1.04 | 1.54 |
| 4JQV | B*18:01 | SELEIKRY | 4XXC | B*18:01 |  | 0.71 | 1.26 |
| 4K7F | A*02:01 | VCWGELMNL | 3H7B | A*02:01 |  | 0.80 | 1.66 |
| 4L3C | A*02:01 | YLLMWITQV | 2BNQ | A*02:01 |  | 1.46 | 2.50 |
| 4MJI | B*51:01 | TAFTIPSI | 1A1N | B*35:01 |  | 1.32 | 2.51 |
| 4N8V | A*11:01 | MLIYSMWGK | 1Q94 | A*11:01 |  | 1.86 | 2.97 |
| 4NNY | A*02:01 | RQASLSISV | 4NO5 | A*02:01 |  | 1.98 | 2.97 |
| 4NO5 | A*02:01 | RQISQDVKL | 4NNY | A*02:01 |  | 1.84 | 2.89 |
| 4NQV | A*01:01 | CTELKLSDY | 4NQX | A*01:01 |  | 0.66 | 1.40 |
| 4NQX | A*01:01 | CTELKLNDY | 4NQV | A*01:01 |  | 0.74 | 1.74 |
| 4NT6 | C*08:01 | GILGFVFTL | 6ULI | C*08:02 |  | 1.85 | 3.70 |
| 4O2E | B*39:01 | SHVAVENAL | 3BXN | B*14:02 |  | 2.45 | 3.90 |
| 4O2F | B*39:01 | HVAVENAL | 3SPV | B*08:01 |  | 3.14 | 4.70 |
| 4QRQ | B*08:01 | HSKKKCDEL | 5WMR | B*08:01 |  | 2.21 | 2.44 |
| 4QRR | B*35:01 | IPSINVHHY | 1A9E | B*35:01 |  | 1.57 | 2.82 |
| 4QRS | B*08:01 | ELKRKMIYM | 4QRT | B*08:01 |  | 2.03 | 2.97 |
| 4QRT | B*08:01 | ELNRKMIYM | 4QRS | B*08:01 |  | 1.04 | 1.72 |
| 4QRU | B*08:01 | ELRRKMMYM | 4QRS | B*08:01 |  | 2.30 | 2.79 |
| 4U1H | B*07:02 | TPQDLNTML | 5EO0 | B*07:02 |  | 0.58 | 1.12 |
| 4U1J | B*42:01 | TPQDLNTML | 4U1M | B*42:01 |  | 1.57 | 3.02 |
| 4U1K | B*07:02 | RPQVPLRPM | 5WMO | B*07:02 |  | 1.76 | 3.59 |
| 4U1M | B*42:01 | RPQVPLRPM | 4U1J | B*42:01 |  | 1.40 | 2.43 |
| 4XXC | B*18:01 | DELEIKAY | 4JQV | B*18:01 |  | 0.58 | 1.29 |
| 5BRZ | A*01:01 | EVDPIGHLY | 3BO8 | A*01:01 |  | 1.48 | 1.90 |
| 5BS0 | A*01:01 | ESDPIVAQY | 5BRZ | A*01:01 |  | 0.60 | 1.28 |
| 5C0D | A*02:01 | AQWGPDPAAA | 3UTQ | A*02:01 |  | 1.20 | 1.98 |
| 5C0E | A*02:01 | YQFGPDFPIA | 5C0I | A*02:01 |  | 0.95 | 2.30 |
| 5C0F | A*02:01 | RQWGPDPAAV | 5C0D | A*02:01 |  | 1.14 | 1.91 |
| 5C0G | A*02:01 | YLGGPDFPTI | 5C0I | A*02:01 |  | 1.05 | 1.65 |
| 5C0I | A*02:01 | RQFGPDFPTI | 5C0E | A*02:01 |  | 0.66 | 1.88 |
| 5C0J | A*02:01 | RQFGPDWIVA | 5C0E | A*02:01 |  | 1.40 | 2.37 |
| 5DEF | B*27:04 | RRKWRRWHL | 2A83 | B*27:05 |  | 2.49 | 4.89 |
| 5E00 | A*02:01 | GVWIRTPPA | 5WSH | A*02:01 |  | 0.82 | 1.77 |
| 5ENW | A*02:01 | GLKEGIPAL | 5FA3 | A*02:01 |  | 0.65 | 1.56 |
| 5EO0 | B*07:02 | RPMTFKGAL | 5EO1 | B*07:02 |  | 0.62 | 1.70 |
| 5EO1 | B*07:02 | RPMTYKGAL | 5EO0 | B*07:02 |  | 0.64 | 1.39 |
| 5EU3 | A*02:01 | YLEPGPVTA | 5EU6 | A*02:01 |  | 1.14 | 2.15 |
| 5EU4 | A*02:01 | YLAPGPVTA | 5EU3 | A*02:01 |  | 1.63 | 2.20 |
| 5EU5 | A*02:01 | YLEPAPVTA | 5EU3 | A*02:01 |  | 1.11 | 2.13 |
| 5EU6 | A*02:01 | YLEPGPVTV | 5EU3 | A*02:01 |  | 1.24 | 2.41 |
| 5F9J | A*02:01 | YLSPIASPL | 7N1A | A*02:01 |  | 1.32 | 2.11 |
|  |  |  | **Template** | |  | **RMSD** | |
| **PDB** | **MHC-I** | **Epitope** | **PDB** | **MHC-I** |  | **Backbone** | **Interface** |
| 5FA3 | A*02:01 | GLLPELPAV | 5ENW | A*02:01 |  | 0.66 | 1.19 |
| 5FDW | A*02:01 | YLSPIASPLL | 5C0G | A*02:01 |  | 1.97 | 2.73 |
| 5GRD | A*11:01 | SSCSSCPLSK | 5WJN | A*11:01 |  | 1.30 | 1.66 |
| 5GSD | A*11:01 | SSCPLSK | 6PBH | A*68:01 |  | 2.65 | 3.13 |
| 5HGA | A*24:02 | RFPLTFGW | 5HGB | A*24:02 |  | 1.24 | 1.78 |
| 5HGB | A*24:02 | RYPLTFGW | 5HGA | A*24:02 |  | 1.43 | 2.00 |
| 5HGD | A*24:02 | RFPLTFGWCF | 3VXN | A*24:02 |  | 1.17 | 1.82 |
| 5HHN | A*02:01 | GILGLVFTL | 1OGA | A*02:01 |  | 0.91 | 1.65 |
| 5HHP | A*02:01 | GILEFVFTL | 1OGA | A*02:01 |  | 0.92 | 3.00 |
| 5HHQ | A*02:01 | GIWGFVFTL | 1OGA | A*02:01 |  | 0.84 | 2.74 |
| 5IB2 | B*27:05 | RRKWRRWHL | 2A83 | B*27:05 |  | 1.16 | 3.61 |
| 5IEK | B*40:02 | REFSKEPEL | 6MT3 | B*18:01 |  | 2.08 | 4.16 |
| 5IM7 | B*58:01 | QASQEVKNW | 5IND | B*58:01 |  | 1.54 | 2.30 |
| 5INC | B*58:01 | QATQEVANW | 5IM7 | B*58:01 |  | 1.87 | 2.71 |
| 5IND | B*58:01 | QASQDVKNW | 5IM7 | B*58:01 |  | 1.30 | 1.88 |
| 5MEO | A*02:01 | ILGKFLHRL | 5MEP | A*02:01 |  | 1.14 | 1.62 |
| 5MEP | A*02:01 | ILGKFLHWL | 4I4W | A*02:01 |  | 0.92 | 2.30 |
| 5MEQ | A*02:01 | ILAKFLHTL | 5MER | A*02:01 |  | 0.75 | 2.19 |
| 5MER | A*02:01 | ILAKFLHEL | 5MEQ | A*02:01 |  | 1.40 | 1.81 |
| 5N1Y | A*02:01 | MVWGPDPLYV | 5C0F | A*02:01 |  | 1.63 | 2.18 |
| 5N6B | A*02:01 | LLWNGPMAV | 6SS8 | A*02:01 |  | 1.13 | 2.10 |
| 5NMH | A*02:01 | SLYNTIATL | 2V2W | A*02:01 |  | 0.75 | 1.32 |
| 5NMK | A*02:01 | SLFNTIAVL | 2V2X | A*02:01 |  | 1.37 | 3.08 |
| 5SWQ | A*02:01 | CVNGSCFTV | 3MRJ | A*02:01 |  | 1.00 | 1.33 |
| 5T6W | B*57:01 | SSTRGISQLW | 5T6X | B*57:01 |  | 1.01 | 1.45 |
| 5T6X | B*57:01 | TSTTSVASSW | 5T6W | B*57:01 |  | 0.95 | 1.25 |
| 5T6Z | B*57:01 | TSTLQEQIGW | 5T70 | B*57:01 |  | 3.58 | 5.11 |
| 5T70 | B*57:01 | TSNLQEQIGW | 5T6Z | B*57:01 |  | 3.17 | 4.66 |
| 5TXS | B*15:01 | AQDIYRASY | 1XR8 | B*15:01 |  | 1.63 | 3.43 |
| 5U98 | B*57:01 | VTTDIQVKV | 5VUF | B*57:01 |  | 2.30 | 3.73 |
| 5V5L | B*58:01 | TSTLQEQIGW | 5T70 | B*57:01 |  | 2.98 | 4.17 |
| 5VGD | C*05:01 | SAEPVPLQL | 6ULI | C*08:02 |  | 1.95 | 3.01 |
| 5VGE | C*07:02 | RYRPGTVAL | 5W6A | C*06:02 |  | 2.36 | 3.11 |
| 5VUD | B*57:01 | LSSPVTKSW | 3VH8 | B*57:01 |  | 0.64 | 0.96 |
| 5VUE | B*57:01 | LTVQVARVW | 5VUF | B*57:01 |  | 0.91 | 1.70 |
| 5VUF | B*57:01 | LTVQVARVY | 5VUE | B*57:01 |  | 1.11 | 1.61 |
| 5VVP | B*57:03 | LSSPVTKSW | 6V2Q | B*57:03 |  | 1.81 | 2.87 |
| 5VWD | B*57:03 | LTVQVARVW | 5VWF | B*57:03 |  | 2.45 | 2.96 |
| 5VWF | B*57:03 | LTVQVARVY | 5VWD | B*57:03 |  | 0.94 | 2.08 |
| 5VWH | B*58:01 | LSSPVTKSW | 5IM7 | B*58:01 |  | 2.21 | 2.82 |
| 5VWJ | B*58:01 | LTVQVARVW | 5VWH | B*58:01 |  | 0.76 | 1.36 |
| 5VZ5 | B*15:01 | AQDIYRASYY | 2AXG | B*35:01 |  | 2.24 | 4.21 |
| 5W69 | C*06:02 | ARFNDLRFV | 5W6A | C*06:02 |  | 0.60 | 2.28 |
| 5W6A | C*06:02 | ARTELYRSL | 5W69 | C*06:02 |  | 1.16 | 2.98 |
| 5WJN | A*11:01 | GTSGSPIINR | 5WKF | A*11:01 |  | 1.18 | 1.78 |
| 5WKF | A*11:01 | GTSGSPIVNR | 5WJN | A*11:01 |  | 0.98 | 1.28 |
| 5WMN | B*07:02 | SPIVPSFDM | 4U1K | B*07:02 |  | 0.55 | 1.04 |
| 5WMO | B*07:02 | RPPIFIRRL | 4U1K | B*07:02 |  | 2.46 | 5.13 |
| 5WMP | B*07:02 | TPRVTGGGAM | 6UJ8 | B*07:02 |  | 2.47 | 3.01 |
| 5WMQ | B*08:01 | ELRSRYWAI | 1M05 | B*08:01 |  | 1.97 | 2.57 |
| 5WMR | B*08:01 | QIKVRVDMV | 4QRQ | B*08:01 |  | 1.58 | 2.26 |
| 5WSH | A*02:01 | GVWIRTPTA | 5E00 | A*02:01 |  | 1.76 | 2.57 |
| 5XOS | B*35:01 | IPLTEEAEL | 6BJ8 | B*35:01 |  | 1.25 | 1.56 |
| 5XS3 | C*06:02 | VRSRRCLRL | 5W6A | C*06:02 |  | 0.80 | 2.31 |
| 6AM5 | A*02:01 | SMLGIGIVPV | 3MRR | A*02:01 |  | 1.21 | 1.53 |
| 6AMU | A*02:01 | MMWDRGLGMM | 5N1Y | A*02:01 |  | 2.64 | 4.85 |
| 6AT9 | A*01:01 | AQDIYRASYY | 1QVO | A*11:01 |  | 2.66 | 3.50 |
| 6BJ8 | B*35:01 | VPLTEDAEL | 5XOS | B*35:01 |  | 0.96 | 1.24 |
| 6D29 | B*57:01 | TSMSFVPRPW | 5T6X | B*57:01 |  | 2.81 | 3.73 |
| 6D2T | B*57:01 | LALLTGVRW | 5VUE | B*57:01 |  | 1.86 | 2.57 |
| 6EWC | A*02:01 | RLSSPLHFV | 1I1Y | A*02:01 |  | 1.63 | 3.05 |
| 6G3J | A*02:01 | MTSAIGILPV | 4JFP | A*02:01 |  | 0.95 | 1.30 |
| 6J1W | A*30:01 | AIFQSSMTK | 6J29 | A*30:03 |  | 1.69 | 2.02 |
| 6JOZ | A*11:01 | ATIGTAMYK | 1Q94 | A*11:01 |  | 1.65 | 1.88 |
| 6MT3 | B*18:01 | FEDLRVLSF | 4QRR | B*35:01 |  | 1.62 | 2.31 |
| 6MT4 | B*37:01 | FEDLRVSSF | 6MT6 | B*37:01 |  | 1.18 | 1.35 |
| 6MT5 | B*37:01 | FEDLRLLSF | 6MT6 | B*37:01 |  | 1.37 | 1.60 |
| 6MT6 | B*37:01 | FEDLRVLSF | 6MT5 | B*37:01 |  | 0.93 | 1.30 |
| 6MTL | B*44:05 | FEDLRVLSF | 1SYV | B*44:05 |  | 1.89 | 3.70 |
|  |  |  | **Template** | |  | **RMSD** | |
| **PDB** | **MHC-I** | **Epitope** | **PDB** | **MHC-I** |  | **Backbone** | **Interface** |
| 6O4Z | A*02:01 | KLVVVAVGV | 6O4Y | A*02:01 |  | 1.31 | 1.64 |
| 6O51 | A*02:01 | YLVVVGAVGV | 6O53 | A*02:01 |  | 1.44 | 1.91 |
| 6O53 | A*02:01 | KLVVVGAVGV | 6O51 | A*02:01 |  | 1.56 | 1.89 |
| 6O9B | A*03:01 | TTAPSLSGK | 6O9C | A*03:01 |  | 0.83 | 1.30 |
| 6O9C | A*03:01 | TTAPFLSGK | 6O9B | A*03:01 |  | 1.01 | 1.40 |
| 6OPD | A*02:01 | ILNAMIVKI | 6PTB | A*02:01 |  | 0.79 | 1.71 |
| 6PBH | A*68:01 | DATALVR | 5GSD | A*11:01 |  | 2.66 | 3.50 |
| 6PTB | A*02:01 | ILNAMIAKI | 6OPD | A*02:01 |  | 1.62 | 2.21 |
| 6PTE | A*02:01 | ILNAMITKI | 6PTB | A*02:01 |  | 1.14 | 1.85 |
| 6PYW | B*27:05 | LRNQSVFNF | 3LV3 | B*27:05 |  | 0.72 | 1.47 |
| 6Q3K | A*02:01 | NLVPMVATV | 3GSR | A*02:01 |  | 0.85 | 1.52 |
| 6R2L | A*02:01 | SLSKILDTV | 5MEQ | A*02:01 |  | 0.98 | 2.03 |
| 6SS7 | A*02:01 | LLWAGPMAV | 5N6B | A*02:01 |  | 1.33 | 2.06 |
| 6SS8 | A*02:01 | LLWNGPIAV | 5N6B | A*02:01 |  | 1.23 | 2.12 |
| 6SS9 | A*02:01 | LLWNGPMHV | 6SSA | A*02:01 |  | 0.94 | 1.64 |
| 6SSA | A*02:01 | LLWNGPMQV | 6SS9 | A*02:01 |  | 0.93 | 1.76 |
| 6TRN | A*02:01 | AVYDGREHTV | 1I4F | A*02:01 |  | 0.86 | 1.23 |
| 6UJ7 | B*07:02 | SPNGTIQNIL | 6UJ8 | B*07:02 |  | 1.43 | 1.70 |
| 6UJ8 | B*07:02 | SPNGTIRNIL | 6UJ7 | B*07:02 |  | 0.83 | 1.50 |
| 6UJO | A*02:06 | KQWLVWLFL | 6UJQ | A*02:06 |  | 1.36 | 2.10 |
| 6UJQ | A*02:06 | KQWLVWLLL | 6UJO | A*02:06 |  | 1.65 | 2.64 |
| 6V2O | B*57:01 | ASLNLPAVSW | 5T6X | B*57:01 |  | 4.55 | 4.11 |
| 6V2P | B*57:03 | ASLNLPAVSW | 5T6X | B*57:01 |  | 3.63 | 4.40 |
| 6V2Q | B*57:03 | LSSPVTKSF | 5VVP | B*57:03 |  | 1.83 | 2.78 |
| 6VMC | A*02:01 | ILDQVPFSV | 1TVH | A*02:01 |  | 1.22 | 1.89 |
| 6VR1 | A*02:01 | HMTEVVRRC | 6VR5 | A*02:01 |  | 1.21 | 2.20 |
| 6VR5 | A*02:01 | HMTEVVRHC | 6VR1 | A*02:01 |  | 0.83 | 1.81 |
| 6XQA | A*24:02 | TYQWVLKNL | 7JYW | A*24:02 |  | 0.72 | 0.96 |
| 6Z9V | A*02:01 | IIGWMWIPV | 6Z9W | A*02:01 |  | 1.73 | 4.35 |
| 6Z9W | A*02:01 | LLGWVFAQV | 6Z9V | A*02:01 |  | 1.50 | 2.62 |
| 7CIQ | B*27:05 | RRFSRSPIRR | 4G8I | B*27:05 |  | 2.39 | 4.06 |
| 7EJL | A*24:02 | QYIKWPWYI | 7EJN | A*24:02 |  | 1.01 | 2.46 |
| 7EJM | A*24:02 | TYIKWPWWV | 7EJL | A*24:02 |  | 1.09 | 2.31 |
| 7EJN | A*24:02 | MYVKWPWYV | 7EJL | A*24:02 |  | 1.15 | 2.27 |
| 7EU2 | A*02:01 | KIADYNYKL | 5HHP | A*02:01 |  | 0.99 | 2.33 |
| 7F4W | A*24:02 | NYNYLYRLF | 7JYW | A*24:02 |  | 1.40 | 2.91 |
| 7JYU | A*24:02 | IYFSPIRVTF | 4F7M | A*24:02 |  | 1.40 | 2.38 |
| 7JYV | A*24:02 | YFSPIRVTF | 2BCK | A*24:02 |  | 1.08 | 2.26 |
| 7JYW | A*24:02 | TYQWIIRNW | 6XQA | A*24:02 |  | 0.79 | 1.41 |
| 7KGO | A*02:01 | ILLNKHIDA | 3FT4 | A*02:01 |  | 1.02 | 1.92 |
| 7KGP | A*02:01 | GMSRIGMEV | 6R2L | A*02:01 |  | 1.35 | 2.50 |
| 7KGQ | A*02:01 | LLLDRLNQL | 5MER | A*02:01 |  | 1.00 | 1.86 |
| 7KGR | A*02:01 | LQLPQGTTL | 3MRD | A*02:01 |  | 1.81 | 3.02 |
| 7KGS | A*02:01 | ALNTPKDHI | 6EWC | A*02:01 |  | 1.59 | 3.25 |
| 7L1B | A*03:01 | AHHGGWTTK | 7L1C | A*03:01 |  | 0.72 | 2.45 |
| 7L1C | A*03:01 | ALHGGWTTK | 7L1B | A*03:01 |  | 0.85 | 1.73 |
| 7LGD | B*07:02 | SPRWYFYYL | 7LGT | B*07:02 |  | 2.15 | 5.28 |
| 7LGT | B*07:02 | SPKLHFYYL | 7LGD | B*07:02 |  | 2.01 | 4.46 |
| 7N1A | A*02:01 | YLQPRTFLL | 5EU6 | A*02:01 |  | 1.32 | 2.61 |
| 7N1E | A*02:01 | RLQSLQTYV | 7KGT | A*02:01 |  | 0.88 | 2.14 |

Supplemental Table 2: FlexPepDock refinement benchmark results summary, including PDB structures used for the benchmark and the PDB structures used as templates.

| **Peptide** | **Sequence** | **Protein** | **Gene** | **NCBI Reference Sequence** | **Start** | **-** | **End** |
| --- | --- | --- | --- | --- | --- | --- | --- |
| P1 | LAGKNLTHI | sodium/glucose cotransporter 2 | Slc5a2 | NP_573517.1 | 303 | - | 311 |
| P2 | SQNKNAKAM | solute carrier family 22 member 2 isoform 1 | Slc22a2 | NP_038695.1 | 296 | - | 304 |
| P3 | FSYKKMTIM | inositol oxygenase | Miox | NP_064361.2 | 66 | - | 74 |
| P4 | SSSKPVEPL | podocin | Nphs2 | NP_569723.1 | 372 | - | 380 |
| P5 | KAMKIIKHI | solute carrier family 22 member 2 isoform 1 | Slc22a2 | NP_038695.1 | 302 | - | 310 |
| P6 | YILKLPLPL | cadherin-16 isoform 1 precursor | Cdh16 | NP_031689.1 | 42 | - | 50 |
| P7 | GSPKQHEVV | solute carrier family 23 member 1 | Slc23a1 | NP_035527.3 | 8 | - | 16 |
| P8 | AAPDHKVVV | cadherin-16 isoform 1 precursor | Cdh16 | NP_031689.1 | 531 | - | 539 |
| P9 | FSLQVKPEI | hepatitis A virus cellular receptor 1 homolog isoform b precursor | Havcr1 | NP_001160104.1 | 124 | - | 132 |
| P10 | MQLPSKVVL | kelch domain-containing protein 7A | Klhdc7a | NP_775603.2 | 17 | - | 25 |
| P11 | SVPTHKSSL | solute carrier organic anion transporter family member 4C1 | Slco4c1 | NP_766246.1 | 255 | - | 263 |
| P12 | MADITKSYL | N-acetyltransferase 8 | Nat8 | NP_075944.1 | 95 | - | 103 |
| P13 | MMKFNKFSL | inositol oxygenase | Miox | NP_064361.2 | 203 | - | 211 |

Supplemental Table 3: List of peptide candidates for *in vitro* and *in vivo* screening.

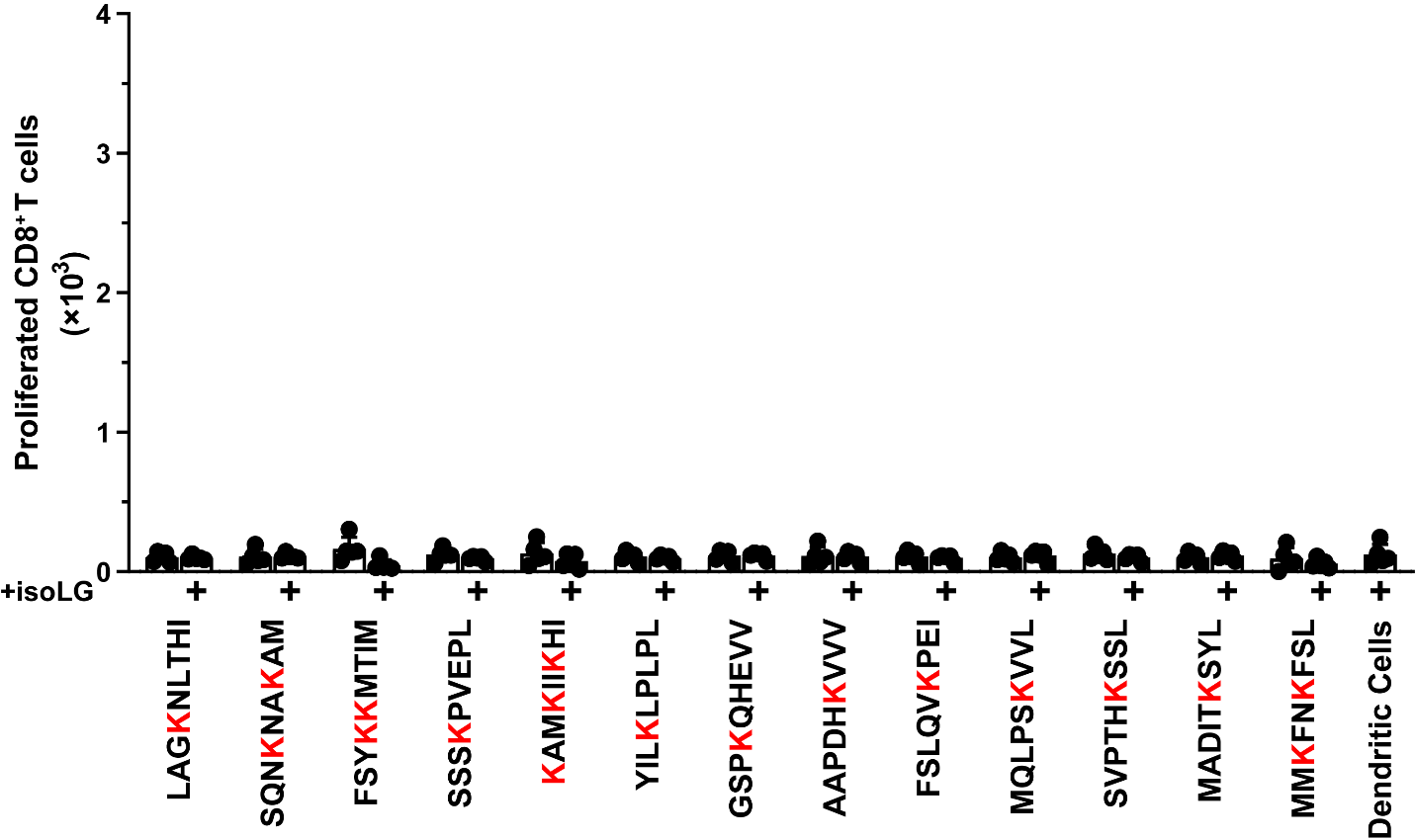

Supplemental Figure 3: T cells isolated from sham treated mice do not proliferate when exposed to isoLG-adducted peptides. n=5, student’s t-test.

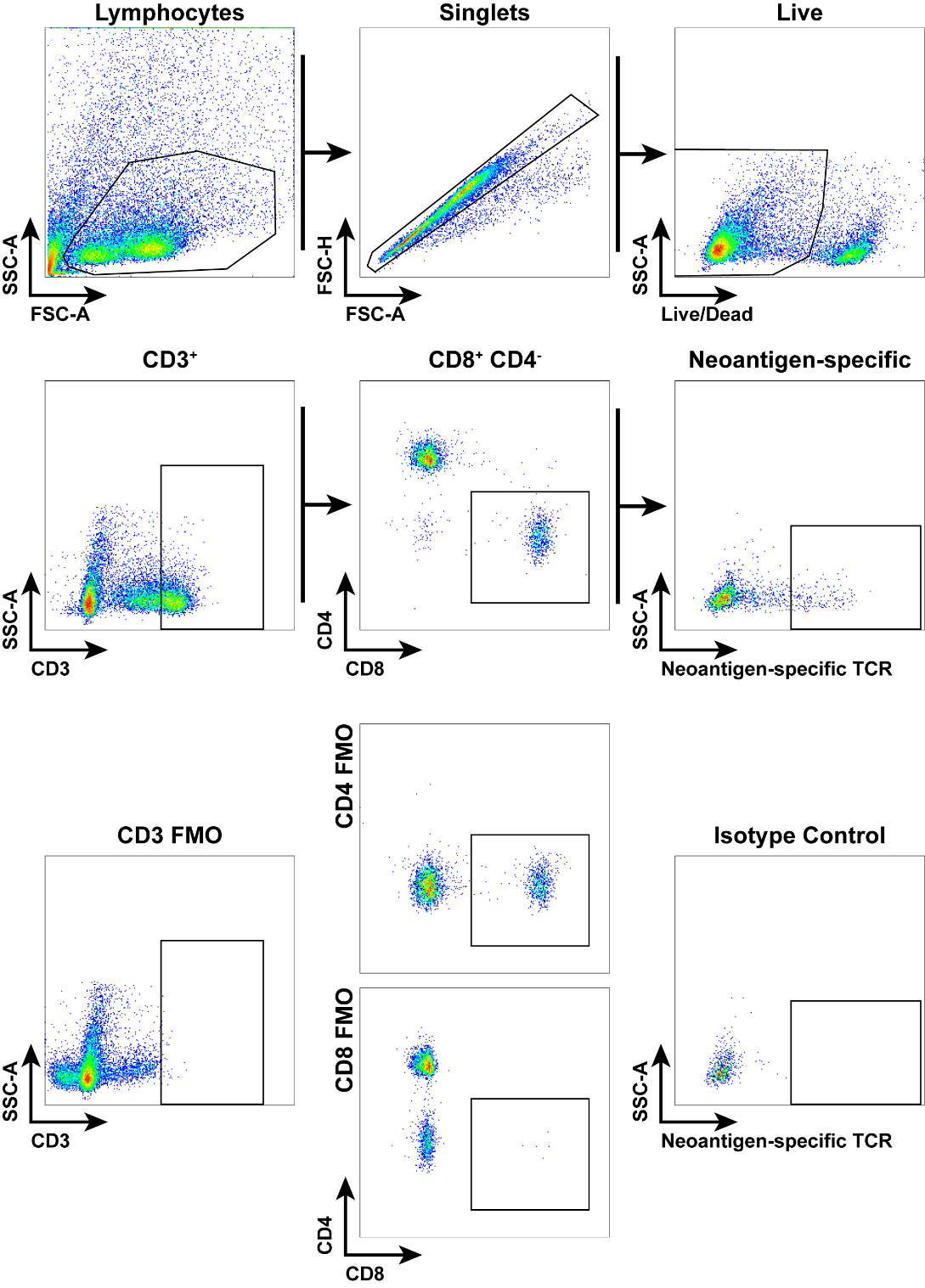

Supplemental Figure 4: Gating strategy and FMOs used for identifying peptide-specific CD8^+^ T cells in peripheral tissues.

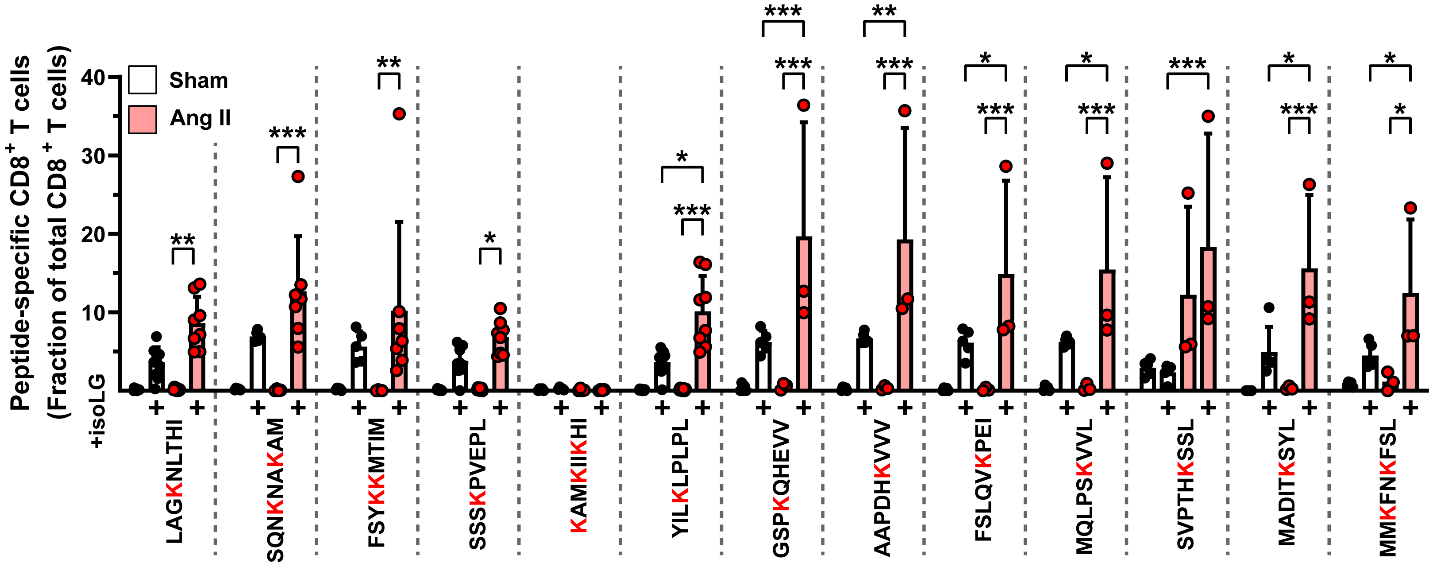

Supplemental Figure 5: The fraction of neoantigen-specific CD8^+^ T cells is increased in the aortas of hypertensive mice. n=3-7, **p*<0.05, ***p*<0.001, ****p*<0.0001, 2-way ANOVA and Holm-Sidak post-hoc test.

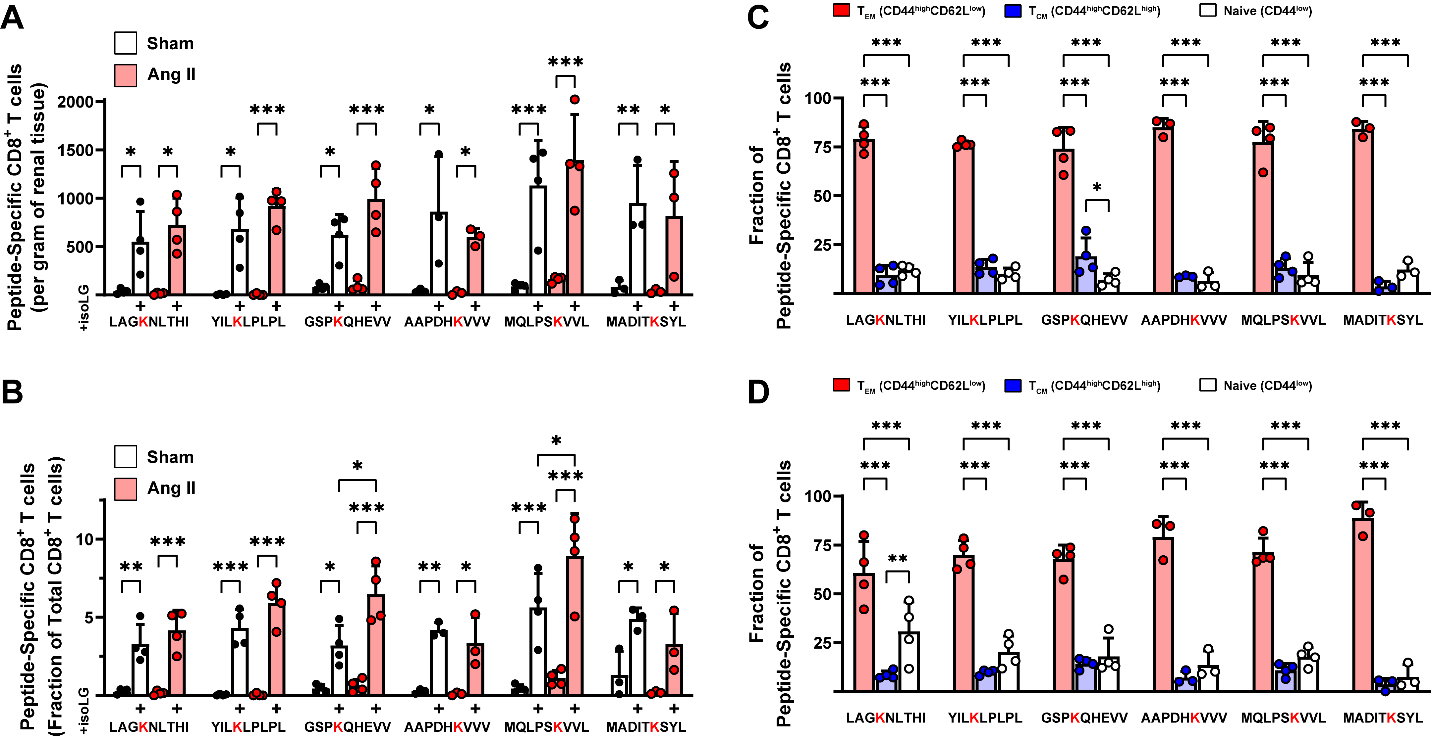

Supplemental Figure 6: Peptide-specific memory CD8^+^ T cells are present in the kidneys of hypertensive mice. (A-B) The fraction, but not total number, of CD8^+^ T cells recognizing two of the six IsoLG-adducted peptides that induce proliferation and identify T cells in the aorta is increased in the kidneys of hypertensive animals compared to sham treated controls (n=3-4, **p*<0.05, ***p*<0.001, ****p*<0.0001, 2-way ANOVA and Holm-Sidak post-hoc). (C) Peptide-specific CD8^+^ T cells in the kidney are predominantly memory effector cells in both (C) sham- and (D) angiotensin II-treated mice. (n=3-4, ***p*<0.001, ****p*<0.0001, 2-way ANOVA and Holm-Sidak post-hoc).

|  |  | Number of alleles selected for screening | | |
| --- | --- | --- | --- | --- |
| Population Name | Size | **HLA-A** | **HLA-B** | **HLA-C** |
| USA NMDP European Caucasian | 1,242,890 | 5 | 8 | 2 |
| USA NMDP African American pop 2 | 416,581 | 4 | 5 | 1 |
| USA NMDP Chinese | 99,672 | 4 | 5 | 1 |
| **Total unique alleles** | | **5** | **10** | **3** |

Supplemental Table 4: Population sizes and HLA alleles screened for IsoLG-presentation. Three of the largest USA NMDP populations available in the HLA allele frequency database were queried, and alleles selected based on frequency of occurrence in that population (>5% phenotype frequency). Duplicate alleles across populations are counted only once in the total selected for screening.

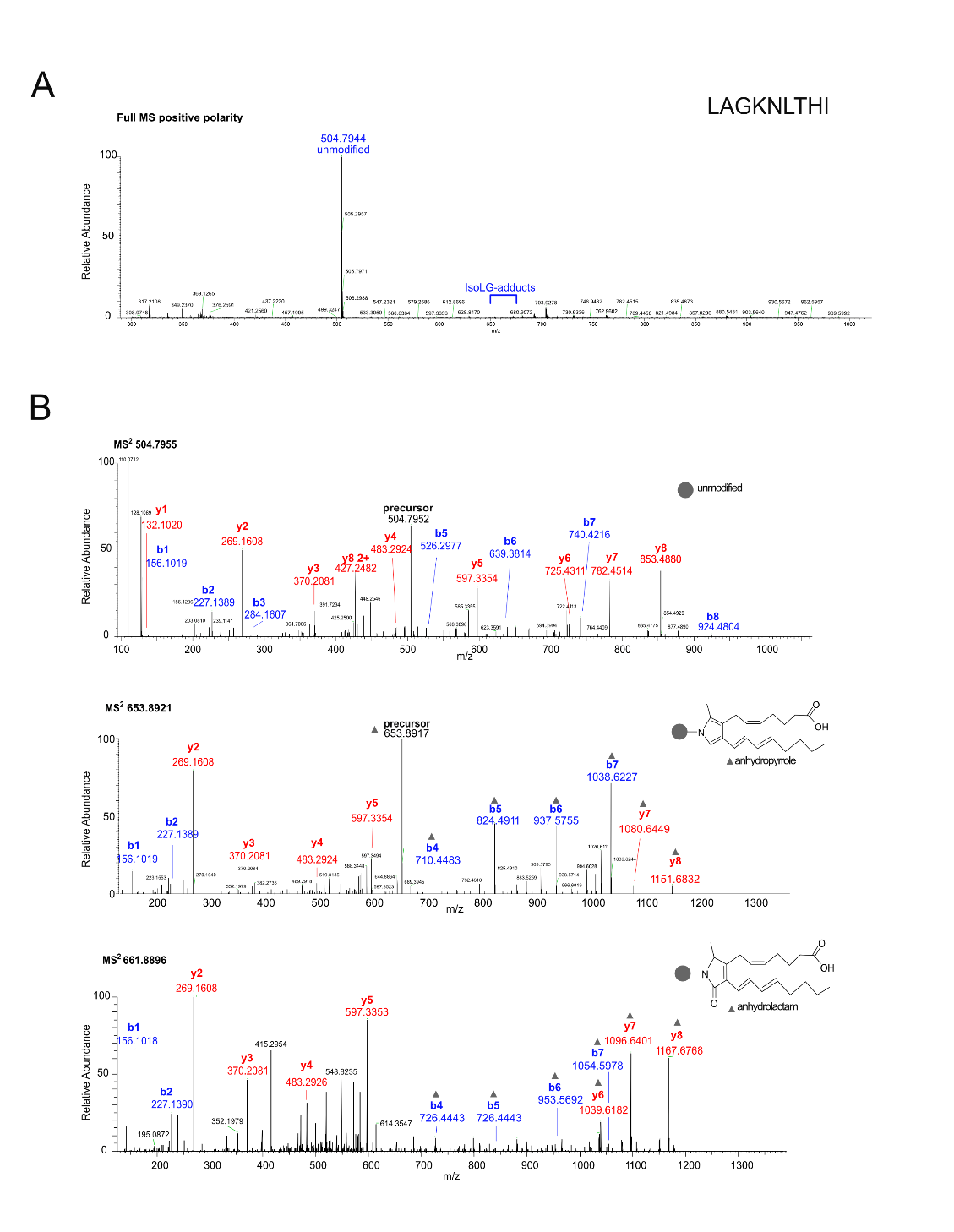

Supplemental Figure 7: Mass spectrometric analysis of the isolevuglandin-E15 (IsoLG) reaction products of the peptide Acetyl-LAGKNLTHI. A) Positive polarity MS of the reaction mixture. IsoLG adducted peptide ions are detectable, but at signal intensities that are two to three orders of magnitude lower than the un-modified peptide. B) MS2 HCD fragmentation spectra of the unmodified peptide, the anhydropyrrole-IsoLG and the anhydrolactam-IsoLG adducted peptide. Series of b- and y- ions are labeled in blue or red, respectively. Fragment ions that carry the IsoLG modification are marked with a solid grey triangle.

Supplemental Figure 8: Targeted mass spectrometric orbitrap parallel reaction monitoring (PRM) analysis of the immunogenic peptide Acetyl-LAGKNLTHI without and after treatment with isolevuglandin-E15. Extracted ion chromatograms are shown for each PRM transition as singly charged y ions. The formation of isobaric IsoLG adduct diastereomers leads to the observation of multiple complex chromatographic peaks for the pyrrole, anhydrolactam, and anhydropyrrole products, simplified structures of which are indicated. The grey circle represents the peptide; its lysine ε-nitrogen atom is included in the adduct structure.

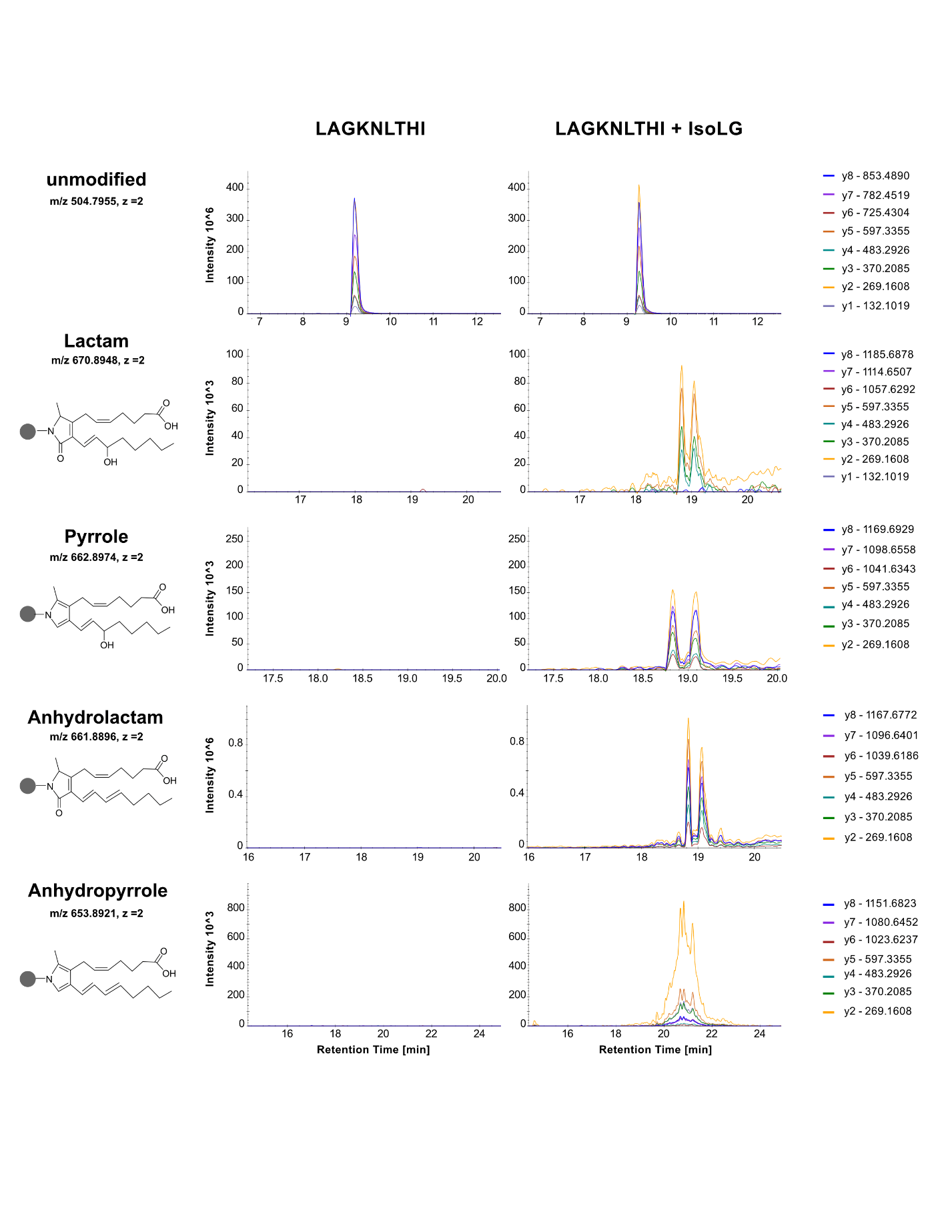

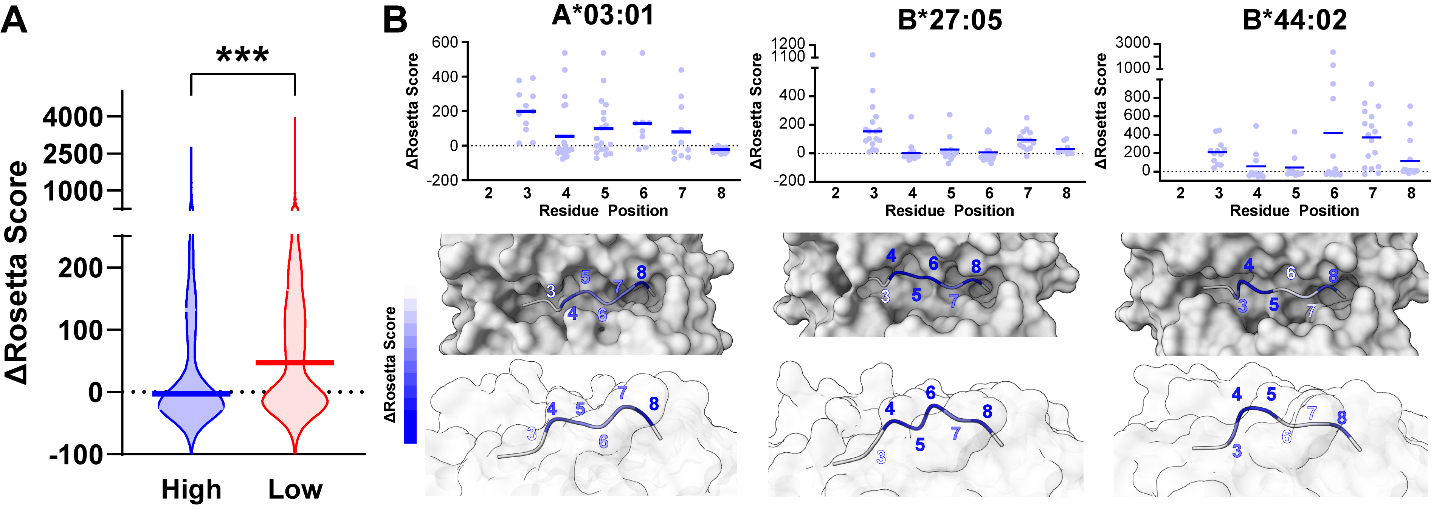

Supplemental Figure 9: Rosetta predicts more favorable energy changes following isoLG adduction for peptides complexed with “high presenting” HLA alleles. (A) Rosetta energy changes are significantly lower (more favorable) following isoLG adduction for peptides complexed with the five high presenting HLA-A and HLA-B alleles compared to the lower-presenting variants (n=472 for High presenters, n=975 for Low presenters, ****p*<0.0001, Mann-Whitney test). (B) Per-residue energy changes following isoLG adduction for peptides bound to high presenters with available crystal structures (top). Residues with more favorable changes are shown in darker blue, and generally correspond to regions of the epitope that jut up and out of the binding cleft (bottom).
